# A Large-Scale Internal Validation Study of Unsupervised Virtual Trichrome Staining Technologies on Non-alcoholic Steatohepatitis Liver Biopsies

**DOI:** 10.1101/2020.07.03.187237

**Authors:** Joshua J. Levy, Nasim Azizgolshani, Michael J. Andersen, Arief Suriawinata, Xiaoying Liu, Mikhail Lisovsky, Bing Ren, Carly Bobak, Brock C. Christensen, Louis J. Vaickus

## Abstract

Non-alcoholic steatohepatitis (NASH) is a fatty liver disease characterized by accumulation of fat in hepatocytes with concurrent inflammation and is associated with morbidity, cirrhosis and liver failure. After extraction of a liver core biopsy, tissue sections are stained with hematoxylin and eosin (H&E) to grade NASH activity, and stained with trichrome to stage fibrosis. Methods to computationally transform one stain into another on digital whole slide images (WSI) can lessen the need for additional physical staining besides H&E, reducing personnel, equipment, and time costs. Generative adversarial networks (GAN) have shown promise for virtual staining of tissue. We conducted a large-scale validation study of the viability of GANs for H&E to trichrome conversion on WSI (n=574). Pathologists were largely unable to distinguish real images from virtual/synthetic images given a set of twelve Turing Tests. We report high correlation between staging of real and virtual stains (ρ = 0.86; 95% CI: 0.84-0.88). Stages assigned to both virtual and real stains correlated similarly with a number of clinical biomarkers and progression to End Stage Liver Disease (Hazard Ratio HR = 2.06, CI 95% 1.36-3.12, P < 0.001 for real stains; HR = 2.02, CI 95% 1.40-2.92, p < 0.001 for virtual stains). Our results demonstrate that virtual trichrome technologies may offer a software solution that can be employed in the clinical setting as a diagnostic decision aid.

## Introduction

Non-alcoholic steatohepatitis (NASH), the most serious condition in the Non-Alcoholic Fatty Liver Disease (NAFLD) spectrum, is a liver disease characterized by serologic markers of hepatocyte injury as well as distinct histological features^1^. NASH is associated with significant morbidity and mortality ^2–4^. Patients with NASH are at increased risk of progressive fibrosis with end-stage liver disease characterized by reduced hepatic synthetic function, portal hypertension, encephalopathy, and malignancy such as hepatocellular carcinoma. Portal hypertension can lead to variceal bleeding, splenomegaly, ascites, and renal injury ^5^. Prevalence of NASH in the United States has been increasing in recent years in parallel with rising levels of obesity and metabolic syndrome^6^. It is predicted to become the leading cause of liver transplantation in the next decade^7^.

Definitive diagnosis of NASH requires tissue biopsy. These biopsy findings can motivate patients to make lifestyle changes (such as improved diet and weight loss) that significantly improve their prognosis. Additionally, appropriate staging can determine if a patient is eligible for routine screening for hepatocellular carcinoma or treatment for varices, ascites, amongst other conditions. Therefore, accurate and timely diagnosis is crucial for patient care. For NASH evaluation, liver biopsies are fixed in formalin, embedded in paraffin and sectioned. Tissue sections are subsequently stained with two principle stain combinations: a routine tissue stain called hematoxylin and eosin (H&E)^8^ for grading of NASH and a trichrome special stain that highlights fibrous tissue blue (or green depending on local staining methods) for staging of fibrosis.^9,10^ Macrovesicular steatosis of hepatocytes, parenchymal inflammatory foci and hepatocyte ballooning are hallmarks of NASH used in grading disease activity while the distribution, extent and interconnectivity of fibrous tissue is used to stage fibrosis.^11^ Maintaining the reagents and technical staff to perform dozens of special chemical stains in a CLIA certified lab is costly and resource intensive^12^. It is thus attractive to computationally “infer” or “translate” a digital WSI of a ubiquitous chemical stain (H&E) to a special stain (trichrome)^13^.

The past 20 years have seen the development of sophisticated computational pipelines that operate on digitized representations of slides, referred to as whole slide images (WSI). Prior to the age of machine learning many of the traditional computational approaches to medical image analysis relied on overly simplistic assumptions as to the shape, color and structure of the tissue features of interest (e.g. nuclei are round, the cytoplasm is pink).^14^ These rigid assumptions often lead to subpar performance on morphometric tasks and fail to capture more informative features that are too esoteric, or high dimensional for pathologists and programmers to effectively capture in rules based programs.^15^ Deep learning approaches, heuristics that utilize artificial neural networks (ANN) and convolutional neural networks (CNN), circumvent this issue in that they are able to find important shapes and patterns for prediction without human specification by capturing and integrating low level features into higher level abstractions^16,17^. Given the advent of these powerful machine learning tools, the Department of Pathology and Laboratory Medicine at Dartmouth Hitchcock Medical Center (DHMC) has fully embraced digital pathology with the caveat that it must leverage advanced computational technologies to provide significant benefits over traditional glass slides.^18–20^ To this end, we have been developing and validating various deep learning technologies in histopathology and designing them to clinical scale^18,21–26^.

Generative image translation techniques transform an image from a source domain A to a target domain B. Recently, these techniques have seen much use in the clinical space, from converting MRI to CT scans to the ability to generate retinal fundus images from blood vessel scans for diabetic retinopathy^27,28^. As such, these technologies present an attractive opportunity in digital histopathology stain translation. The most promising of these methodologies rely on the use of generative adversarial networks (GAN), which generate realistic images from noise (for the purpose of this study, an input signal) using two ANNs; one, the generator, which “generates” the images, and another, the discriminator, which decides whether an incoming image is real or virtual/synthetic^29^. The quality of images created by GANs improves as the generator “learns” to fool the discriminator through an iterative process of trial and error. GANs have been applied in histopathology with success, including for stain translation from autofluorescence imaging^30^, to removal of technical artifacts, as well as for nucleus detection and augmentation of deep learning datasets with synthetic images to improve prediction accuracy.^13,30–42^

Our initial preliminary evaluation of technologies for inferring trichrome stains from H&E featured a small dataset of images to test the viability of utilizing such techniques at our institution.^23^ Here, we perform a full-scale internal validation study of inferring trichrome stains from H&E at our institution for near-term incorporation into our clinical workflow. We describe an image translation model for the purpose of converting H&E stains into trichrome stains, validate the technology through comparison of the correlation in staging real versus virtual stains with a preliminary non-inferiority cutoff, and highlight technical improvements that demonstrate its potential readiness for clinical deployment.

## Methods

### Study Population

Our study cohort consisted of 273 individuals with biopsy proven NASH diagnosis from Dartmouth Hitchcock Medical Center in Lebanon, New Hampshire. To identify patient cases for our study, we performed a keyword search through a Cerner database for the words “trichrome” OR “steatohepatitis” OR “steato” OR “macrovesicular”, OR “steatosis”, OR “NASH” OR “alcoholic” over all pathology cases from July 2007 to March 2019. Identified cases were then individually checked to make sure the case was actually diagnosed as steatosis / steatohepatitis of the non-alcoholic type. All patients had paired H&E and trichrome sections (574 total WSI; 14 patients had repeat biopsies; 21 of the 287 liver biopsies were wedge biopsies), taken at DHMC. Retrospective demographic data, serologic markers, and symptoms of end stage liver disease were collected from patient charts. We chose to collect data on patient characteristics that have been shown to correlate with fibrosis score and prognosis such as BMI ^43^, status of having a diagnosis of impaired fasting glucose, and diabetes mellitus^44^. Additionally, in order to address potential discrepancies between scores of traditionally stained tissues and their virtual counterparts, we collected data on non-invasive markers of hepatic fibrosis that are widely used. AST and ALT are well known independent predictors of NASH induced liver fibrosis^45^. Markers of reduced hepatic function such as INR, albumin and downstream effects of portal hypertension such as hyponatremia, thrombocytopenia, creatinine levels were also recorded. Sodium levels were capped at 137 milliequivalents per liter as values greater than that are not found to be predictive of disease severity and this is the value used to calculate the MELD score ^46^. We collected the serologic data from the time of diagnosis and last clinical encounter.

Lastly, we collected serologic markers in order to calculate MELD and Fib4 scores which have been shown to correlate with outcomes of steatohepatitis. Fib4 combines platelet count, ALT, AST and age to calculate a score which has been shown to have good predictive accuracy for advanced fibrosis in NASH ^47–49^. The MELD score is a prospectively developed and validated measure of liver disease severity commonly used by United Network for Organ Sharing (UNOS) to assess the need for transplantation ^50^. One-Way Analysis of Variance (ANOVA) and Chi-Squared statistical tests were performed to compare these demographic statistics between the median fibrosis stages assigned to the real slides.

### Biopsy Collection and Digitization

Liver tissue from our cohort of NASH patients were obtained by core needle biopsy, fine needle biopsy, and wedge resection. Samples were fixed and embedded in paraffin blocks and 5-micron sections were stained with H&E and trichrome. Slides were scanned using the Leica Aperio-AT2 (Leica Microsystems, Buffalo Grove, IL) scanner at 20x magnification, stored in SVS image format (JPEG compression at 70% quality), from which they were extracted and converted to NPY (uncompressed unsigned 8-bit RGB numpy array) format via our in house pipeline, *PathFlowAI*. ^18^

We utilized 20 H&E/Trichrome WSI pairs for training the deep learning model, spanning a representative sample of fibrosis stages. WSI are large images that can measure hundreds of thousands of pixels in any given spatial dimension. As such, they are currently too big to fit into memory for a deep learning analysis and must be broken into smaller sub-images. In this case, a whitespace detector contained within the *PathFlowAI* software framework, was used to extract 33,456 non-overlapping 512-pixel subimages that contained up to 8% of white space, correspondent to the 40 training WSI.

### Experimental Design

We utilized a CycleGAN architecture because of its clear advantages in translating unpaired images ^51,52^. More details and justification on model selection may be found in the supplementary section “Model Selection”. We trained a CycleGAN model on the 33,456 subimages from 20 paired WSI, randomly cropping 256-pixel subimages from each of these 512-pixel tiles to augment the amount of usable data. GANs can train unpredictably and thus may not have an objective to converge. Instead, the user must visually inspect images during the training process to verify training iterations where the model is producing highly realistic images. However, judging the quality of the any given subimage was not sufficient to decide whether the translation technique could produce both highly realistic and clinically actionable trichrome WSI. Therefore, we built into the currently existing CycleGAN framework the ability to test the latest iteration of the model on an entire WSI by dynamically extracting and translating overlapping subimages from the H&E WSI, then blending overlapped areas as they were used to construct the virtual trichrome. This ensured maximum realism and reduced the appearance of obvious “tiling artifact". During training, we selected two representative H&E/Trichrome pairs (demonstrating different distributions and intensities of trichrome staining) to dynamically translate at various points during model training. A pathologist then visually compared the predicted trichrome WSIs to the actual trichrome WSIs at a variety of points during model training (every 5000 training batch iterations; Figure 1a). After training the model across 200,000 batch iterations (Figure 1b), a pathologist selected the top model through visual comparison of the real and virtual trichrome WSI generated for these two images (Figure 1c). We applied the chosen CycleGAN model to virtually trichrome stain 287 H&E WSI using an inhouse pipeline that encases the selected model, performs the translation and exports WSI into formats that pathologists can read through open-source applications like QuPath or ASAP ^53^. Our in-house pipeline also deploys the slides onto a password-protected image server that allows for tissue-staging from any private web browser through the use of openseadragon^54^ (Figure 1d). Details about the software implementation for training and slide viewing may be found in the Supplementary Materials (section “Software Implementation”).

**Figure 1:**
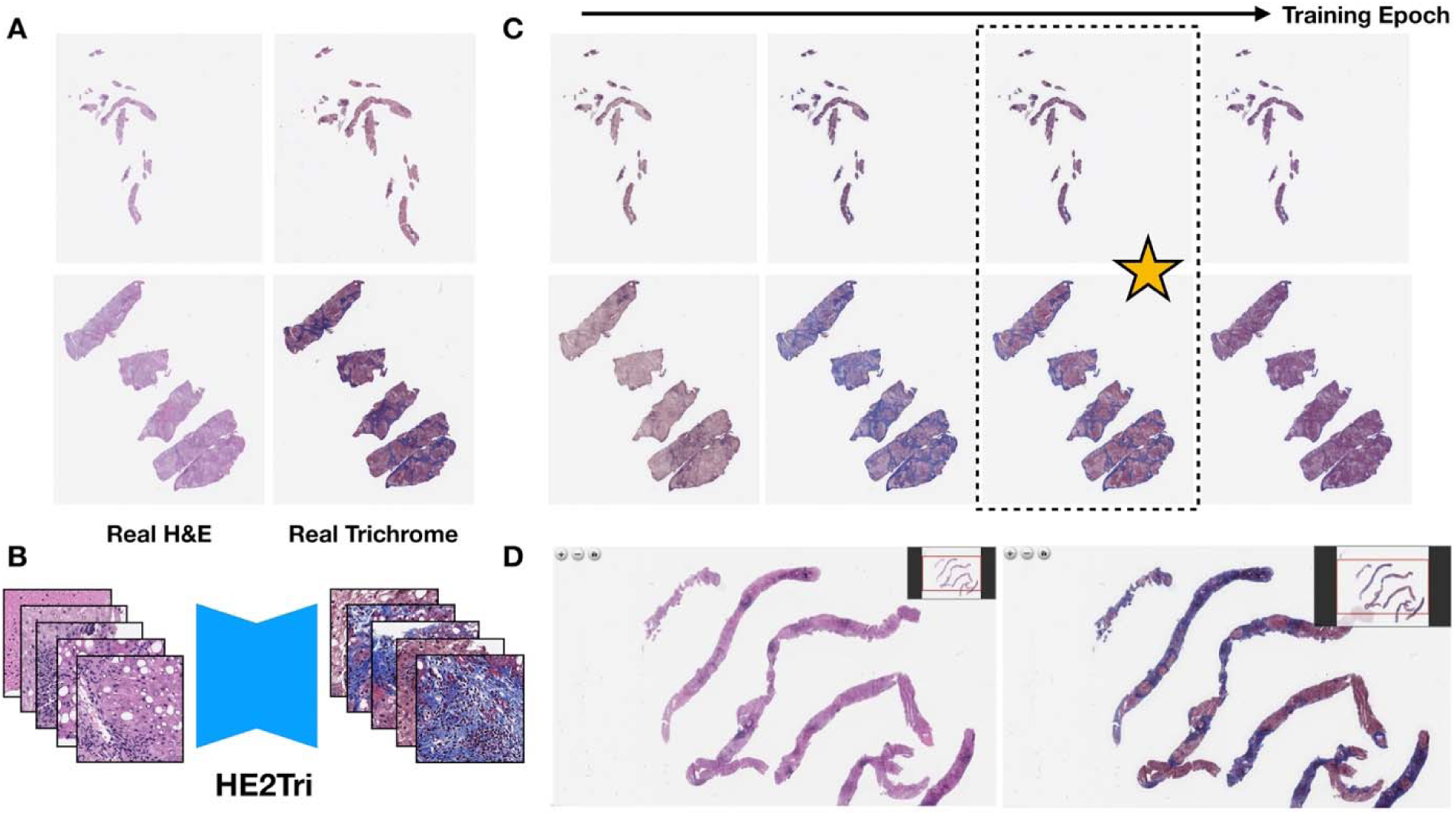
Overview of Workflow: a) Original H&E and trichrome stained slides; b) H&E patches are converted into trichrome patches using CycleGAN; c) Entire trichrome WSI are generated at various training iterations; ideal model is selected; d) Final model is deployed and real time generated/virtual WSI (right) are presented next to original WSI (left) in OpenSeaDragon-based image server for pathologist review ^54^

Four board-certified pathologists (all of whom regularly stage fibrosis in liver biopsies on the gastrointestinal subspecialty service) independently staged the virtual and real trichrome stains via the NASH CRN system ^55^. The inspection of virtual and real images was separated by a period of at least two weeks to reduce the risk of the pathologist recognizing previously staged counterparts in the virtual image set. The real image set was also numbered and ordered differently from the virtual set. All pathologists staged separately the virtual and real stains for each of the slides, (8 total assessments per slide; 4 assessments for real, 4 for virtual; one pathologist was not able to determine stage for two slides; 287 slides; 8*287-2=2294 observations). We also acquired a second set of stages for the real slides as further validation over the course of two months following another washout period of at least one month (separate set of n=1145 observations, a stage could not be determined for three images; total set of n=3439 observations, correspondent to 2291 pairs of assigned virtual and real stages). Images were staged using the NASH CRN system, a five-point ordinal scale with F0 indicating no fibrosis, F1 as portal fibrosis without septa, F2 as few septa, F3 as numerous septa, bridging fibrosis without cirrhosis, and F4 as cirrhosis ^55,56^. Image inspection was conducted using both ASAP Slide V (after a TIFF file export) and openseadragon; the usage of either system by an individual pathologist was dependent on computational limitations (e.g. some of the pathologists’ work and personal computers could not open large images due to hardware limitations) and familiarity with the software. We felt the selection of viewing software would have negligible impact on the ability to stage as both offer a nearly identical viewing experience and have identical controls.

### Assessment of Visual Quality of the Tissue with a Turing Test

We solicited feedback on the visual quality of the real and virtual stain slides in the form of optional pathologist notes compiled in a table by the pathologists as they screened the virtual slides. Given the highly subjective and sporadic nature of solicited feedback, we found that these recordings did not quantitatively communicate the extent of the presence of artifacts and their impact on liver fibrosis staging. To quantitatively assess visual quality of our stains, we conducted multiple Turing Tests, the most common method to assess the quality of a translated image using GANs. Typical, tests are conducted by showing pathologists images of real and virtual subimages, randomly sampled across the real and virtual stained slides. Each pathologist assesses whether the image is real or virtual, the results of which are tabulated against the ground truth to yield a contingency table. A Chi-Squared or Fischer’s Exact Test assesses whether the virtual images may be easily separated from the real images, and vice versa. A Turing Test passes if the corresponding statistical test (Fischer’s Exact) fails to reject the null hypothesis. We presented each pathologist with a random set of 100 real and 100 virtual images for three separate subimage sizes: 256×256, 512×512, and 1024×1024 pixels, where larger images are more likely to contain disqualifying tissue artifacts (200 images per subimage size* 3 subimage sizes * 4 pathologists = 2400 observations). We conducted a total of twelve Turing Tests, four tests (one per pathologist) for each subimage size to assess the visual quality of the virtual stain.

### Quantitative Assessment of Concordance

To validate our technology, we aimed to design a preliminary non-inferiority test to demonstrate concordance between pathologist staging of the real trichromes and the corresponding virtual trichromes, which could address many of the challenges associated with assessment using the NASH CRN system (see supplementary materials section “Challenges with Assessment via NASH CRN System and Justification for Statistical Modeling Approach” and “Treatment of Pathologist Staging Uncertainty for Study Evaluation”). As such, we designed a modeling approach that would attempt to account for ordinal outcome measurements ^57^, pathologist bias (eg. was the pathologist predisposed to increasing or decreasing the assigned virtual stage given the real stage), memorization from repeated testing of real stages, and repeated measurements within case, while establishing a cutoff value that was commensurate with the difficulty of the task at hand (non-inferiority of a GAN technology). To account for these considerations, we correlated real stage to virtual stage using a multi-level ordinal regression model with the following functional specification:

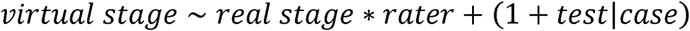

Real stage was modeled as a monotonic effect ^58^. The interaction term, *real stage * rater*, accounts for pathologist/rater bias. The random slope, *(1 + test|case)*, accounts for memorization between successive tests of real stages given repeated measurements per case. Virtual stage was modeled from these predictors using ordinal logistic regression with a cumulative link model, which assumes that fibrosis stages represent “cut points” in a latent distribution of the true underlying continuous fibrosis progression ^59–61^. To handle uncertainty in pathologist assigned stage (521 observations, or 15% of the dataset featured reported interval measurements to convey uncertainty, eg. F2-F3; impacts 67% of the slides), we separately considered the upper bound (eg. F3) and the lower bound (eg. F2) of the measurement interval. To account for high interrater discordance, we separately considered the modeling of concordance in the scenario where we retained cases with high agreement of real stain staging between the pathologists (as quantified by an interquartile range (IQR) in real stages less than or equal to 0.3). In each case, parameters were estimated using a hierarchical Bayesian framework, which utilized Markov Chain Monte Carlo procedures with the R package *brms*. We integrated over the predictive posterior to obtain bias adjusted ordinal estimates of fibrosis stage, which were correlated to virtual stage using Spearman’s Correlation Coefficient. A 95% confidence interval for the correlation coefficient was estimated via a 10,000 sample non-parametric bootstrap and compared to a preliminary non-inferiority cutoff of 0.9^62^. Additional information on the statistical modeling procedures, convergence plots and treatment of pathologist uncertainty in staging may be found in the supplementary material, under sections “Treatment of Pathologist Staging Uncertainty for Study Evaluation” and “Statistical Methods and Results”.

We tested for inter-rater variability through calculation of Spearman’s Correlation Coefficient between pairs of pathologists for the set of virtual stages and both sets of real stages. We separately tested for test-retest reliability ^63^ of real staging (comparing results from the first real stain stages to the second set) using a 10,000-sample non-parametric bootstrap of spearman’s correlation coefficient after acquiring two sets of real stages per slide from each pathologist.

The CycleGAN model was used to make patch level predictions and was thus not tied to staging being done on the WSI-level. In this spirit (level of analysis), and with regards to preliminary results that demonstrated that including the training slides in the evaluation set had negligible impact on concordance, we utilized the entire set of slides for the correlation analyses.

After performing a correlation analysis, which assessed overall agreement across a wide range of stages, we wanted to understand whether pathologists could accurately assess virtual slides for the patients with the highest potential for progression to cirrhosis. We assessed sensitivity for staging advanced fibrosis (defined as median fibrosis stage equal to F4) in slides that were staged in complete pathologist agreement for real stains by recording proportion of times that each pathologist staged a given virtual trichrome as advanced fibrosis. We averaged these accuracy scores to yield a final agreement measure for advanced fibrosis. We then assessed for agreement for the presence or absence of advanced fibrosis for the entire set of pathologist-determined real and virtual fibrosis stages (dichotomized median real and virtual stage by strict cutoff *F = 4*). Cohen’s Kappa statistic was used as the measure of concordance for this assessment.

### Assessing Relationships with Clinical Endpoints

In order to further test the accuracy of the virtual trichrome stain, we aimed to assess if the staging obtained by virtual and real stains were not only correlated with each other but with serologic markers of advanced fibrosis. These markers provide us with a means of assessing discordance between the virtual and real staging means against quantitative serological data which is known to be correlated with fibrosis / NASH severity. We utilized Spearman’s Correlation Coefficient to correlate known serological markers with median virtual staging, real staging, the residual between the real and virtual assigned stage, and the variation in pathologist staging of the tissue (as defined by the IQR). For the sodium biomarker, we capped the maximum value at 137 and any value greater was marked as 137 as is the protocol in calculating MELD scores^46^. We also correlated fibrosis staging with the AST/ALT ratio, of which an AST/ALT ratio greater than one is associated with bridging fibrosis ^45^.

We additionally used Kaplan-Meier survival analysis to determine the relationship between virtual and real fibrosis stage with development of end stage liver disease. Our model adjusted for age and BMI. An event was defined as the occurrence of ascites, hospitalization due to liver disease, encephalopathy, variceal bleed, evidence of varices with imaging, a diagnosis of hepatocellular carcinoma, dialysis, or the prescription of non-specific beta blocker propranolol as a prophylaxis for esophageal variceal bleed during follow up. Multivariate cox proportional hazard models adjusting for age and sex were also used. As these are markers of late stage liver disease, we also dichotomized our cohort into those with a virtual fibrosis stage greater than or equal to F3.5 and less than F3.5 and re-ran our analyses. For the serological and survival analysis, we modeled median stage as a continuous variable, which is considered to be an acceptable practice for the treatment of ordinal independent variables by maintaining the parsimony of the relationship with the outcome while reducing degrees of freedom ^64,65^.

## Results

### Results Overview

We conducted a large-scale internal validation study of H&E to trichrome conversion by training a CycleGAN model on subimages from 20 paired WSI. Then, we converted our entire set of 287 H&E stained WSI into trichrome stained WSI; when processed in series, it took an average of 5 minutes to convert an entire H&E WSI into a virtual trichrome stained WSI. While other studies have chosen to assess the quality of the stain as the means of assessing the synthetic images, which is only tangentially related to the true utility of the virtual stain, we believe that the best way to assess the viability of the approach is whether pathologists can still assign the same fibrosis stage as if they were examining the true trichrome stain on an adjacent tissue section (after accounting for bias/interrater variability). Assessments of virtual trichrome stains should also correlate with actual clinical endpoints (eg. progression to cirrhosis, serological markers of liver injury and failure) in the same manner as the real stain. Our results indicate that the CycleGAN model we trained is capable of generating virtual trichrome stains from H&E which may be sufficient for clinical use pending additional validation.

### Patient Demographics

We tabulated patient demographic information (age, sex, BMI, various serological markers such as Fib4, AST, ALT) from our study cohort, which is summarized in Supplementary Table 1. Calculation of chi-squared statistics for categorical demographics and ANOVA tests for continuous outcomes demonstrated statistically significant differences in some but not all of the serological markers across the mean Fibrosis stages for the real trichrome slides.

The mean age of our cohort was 53, ranging from 8-81. Consistent with previous studies, our cohort had an elevated BMI with a mean of 34 ^43^. Table 1 displays characteristics of our study population stratified by fibrosis score obtained by inspection of the real trichrome. The majority (60%) of our patient cohort had significant fibrosis with a NASH CRN score greater than or equal to F2.

### Pathologist Staging and Histological Inspection

When comparing pairs of real and virtual trichrome images there was generally excellent visual correspondence and for these cases it was difficult to appreciate a difference at low magnification (Supplementary Figures 1-5). For stage 0 fibrosis (figure 2a) the virtual trichrome demonstrates hazy, serum-like staining in the periphery of the tissue and slight understaining of the wall of the central vein. In stage 1 fibrosis (figure 2b), the virtual stain shows patterns consistent with chicken wire fibrosis (due to perisinusoidal scarring) although to a lesser extent than in the analogous field in the real trichrome stained slides. In stage 2 fibrosis (figure 2c) the virtual stain shows similar levels of portal and chicken-wire fibrosis with some aberrant edge staining. In stage 3 fibrosis (figure 2d) the virtual stain demonstrates a nearly identical distribution of fibrosis as compared to the real image with slight edge-staining. In stage 4 fibrosis (figure 2e) the virtual fibrosis staining pattern is virtually indistinguishable from the real trichrome. During pathologist review of the virtual trichrome WSI, the most frequent artifacts encountered were aberrant edge staining in the absence of capsule, and patchy overstaining / understaining. In some cases, the virtual image was subjectively overstained (figure 3a). In this example, a low magnification inspection demonstrates areas of both overstaining and understaining (large portal area). At higher magnification there is obvious portal, parenchymal and edge overstaining in the virtual stain. An example of understaining of the virtual image (figure 3b) is apparent in both the portal and parenchymal areas.

**Figure 2:**
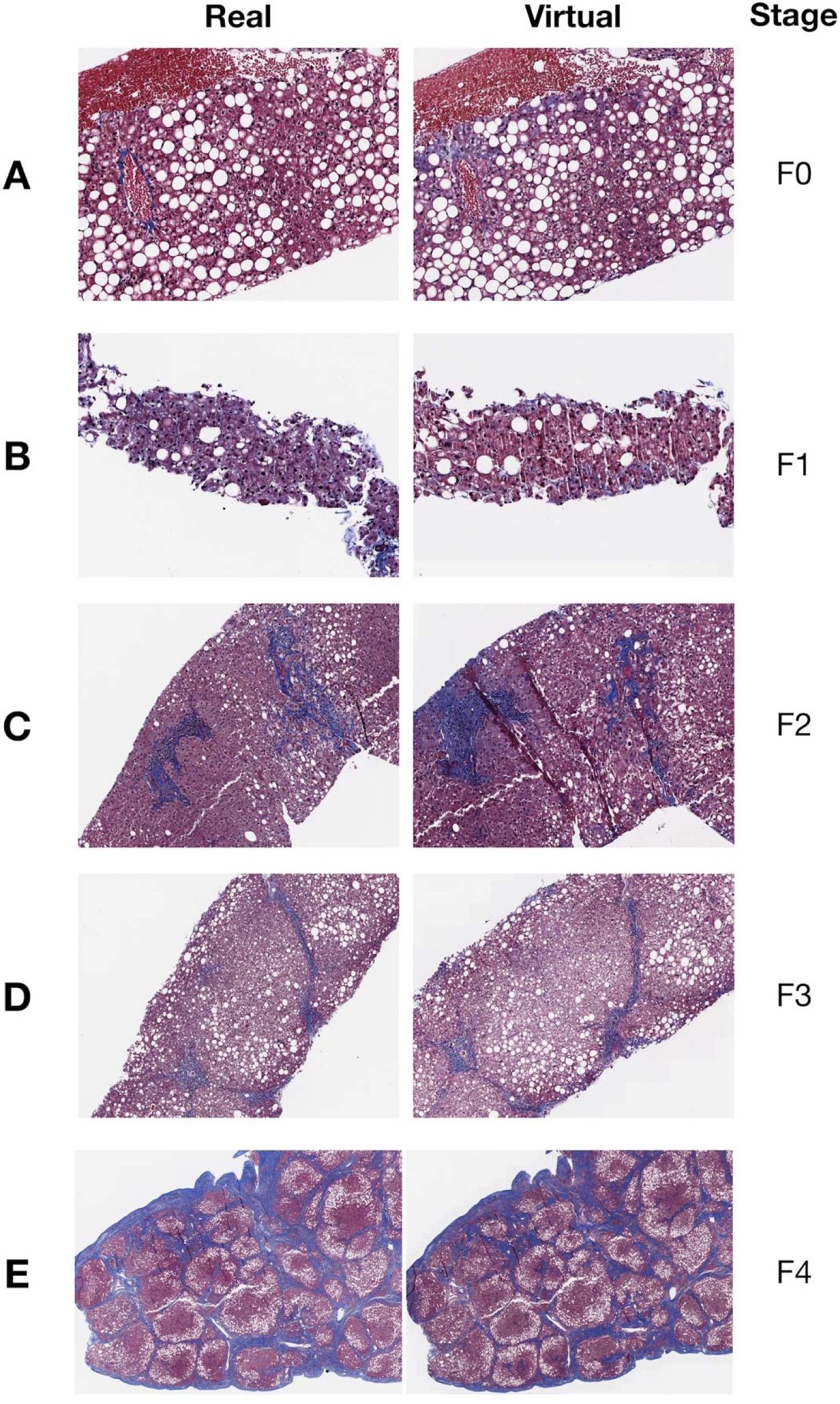
Select Regions of WSI across different accepted Fibrosis stages by: a) Real trichrome, b) Virtual trichrome stains; Entire WSI (including H&E stain) are found in Supplementary Figures 1-5

**Figure 3:**
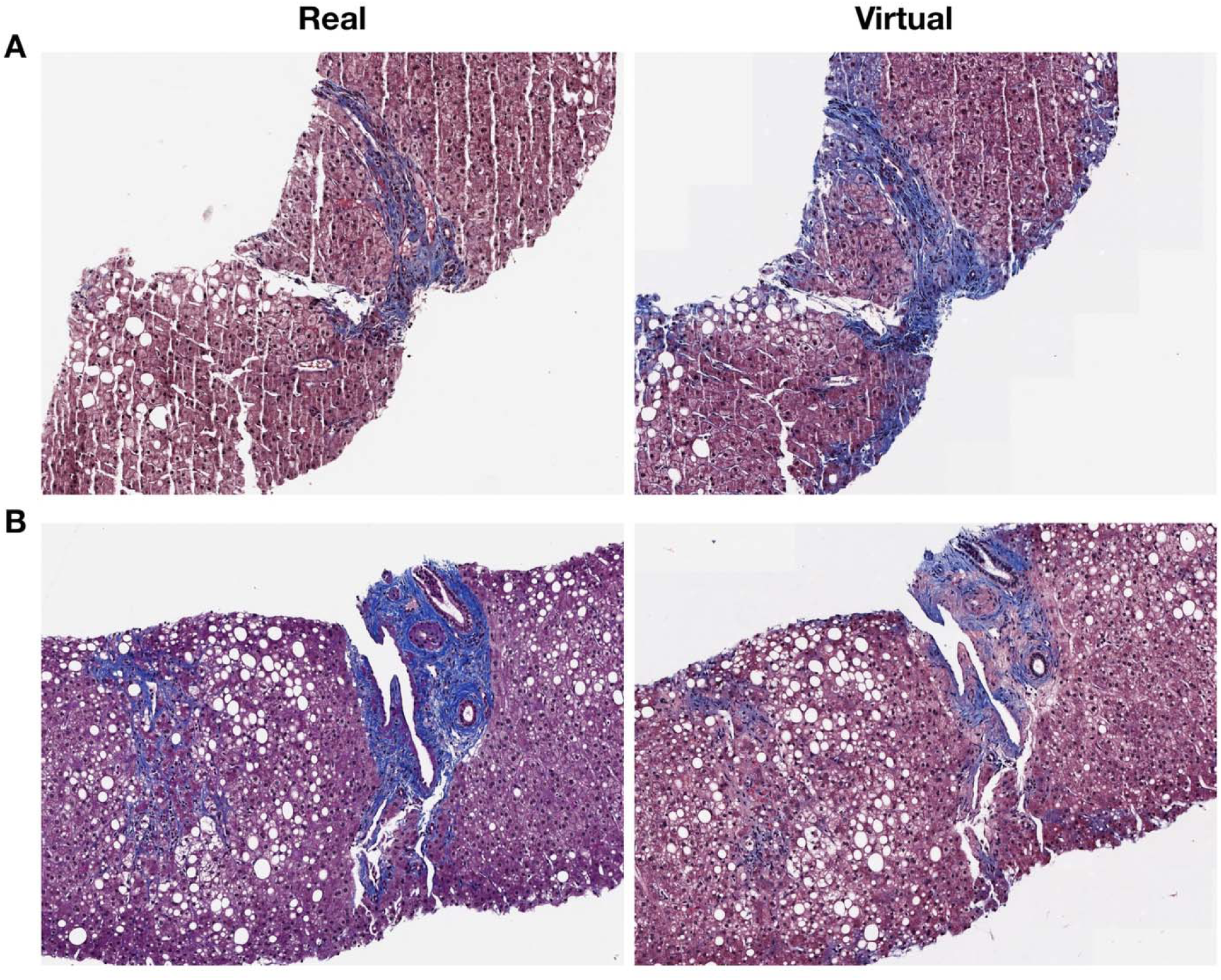
Examples of Virtual Staining Artifacts: a) Overstaining; b) Understaining

### Quantitative Assessment of Stain Quality via Turing Test

While there were additional understaining/overstaining artifacts in addition to artifacts related to biopsy procurement (Supplementary Table 2), the quality of the majority of the virtual stains was excellent. To qualify this assessment, pathologists were unable to distinguish a virtually stained subimage from a real subimage in nine out of the twelve Turing Tests (Supplementary Figures 6-7; Supplementary Table 3). None of the four pathologists were able to distinguish real from virtual subimages for images of size 256×256 (p=0.55, p= 0.06, p=0.64, p=0.07); only one pathologist could distinguish images of size 512×512 (p=0.01, p=0.30, p=0.44, p=0.56); two out of the four pathologists were able to distinguish images of size 1024×1024 (p<0.001, p=0.16, p=0.14, p=0.004), which indicates that larger image crops contained a higher likelihood for disqualifying image artifacts. Overall, the results indicate high micro and macro architectural quality when inspecting various slide subregions.

However, we felt that the common artifacts would have negligible impact on the ability for the pathologist to stage the slide. Thus, we sought to quantify the concordance between the clinical staging of the real and virtual slides to assess the true viability of staging virtual stains while accounting for rater subjectivity.

### Correlation between Real and Virtual Stain Stage

When assessing all slides and considering the upper bound of pathologist measurements for ambiguous assignments for real and virtual stain stages (eg. upstaging F2-3 to an F3) (Figure 4a,c-d), the correlation between bias-adjusted real and virtual stage was 0.81 (95% CI: 0.79-0.83). When considering the cases with low interrater disagreement of the real stages, we obtained a bias-adjusted correlation of 0.86 (95% CI: 0.84-0.88), which fell slightly short of the preliminary non-inferiority correlation cutoff of 0.9. Complete concordance statistics can be found in the supplementary material (section “Statistical Methods and Results”; Supplementary Figure 8, Supplementary Table 4).

**Figure 4:**
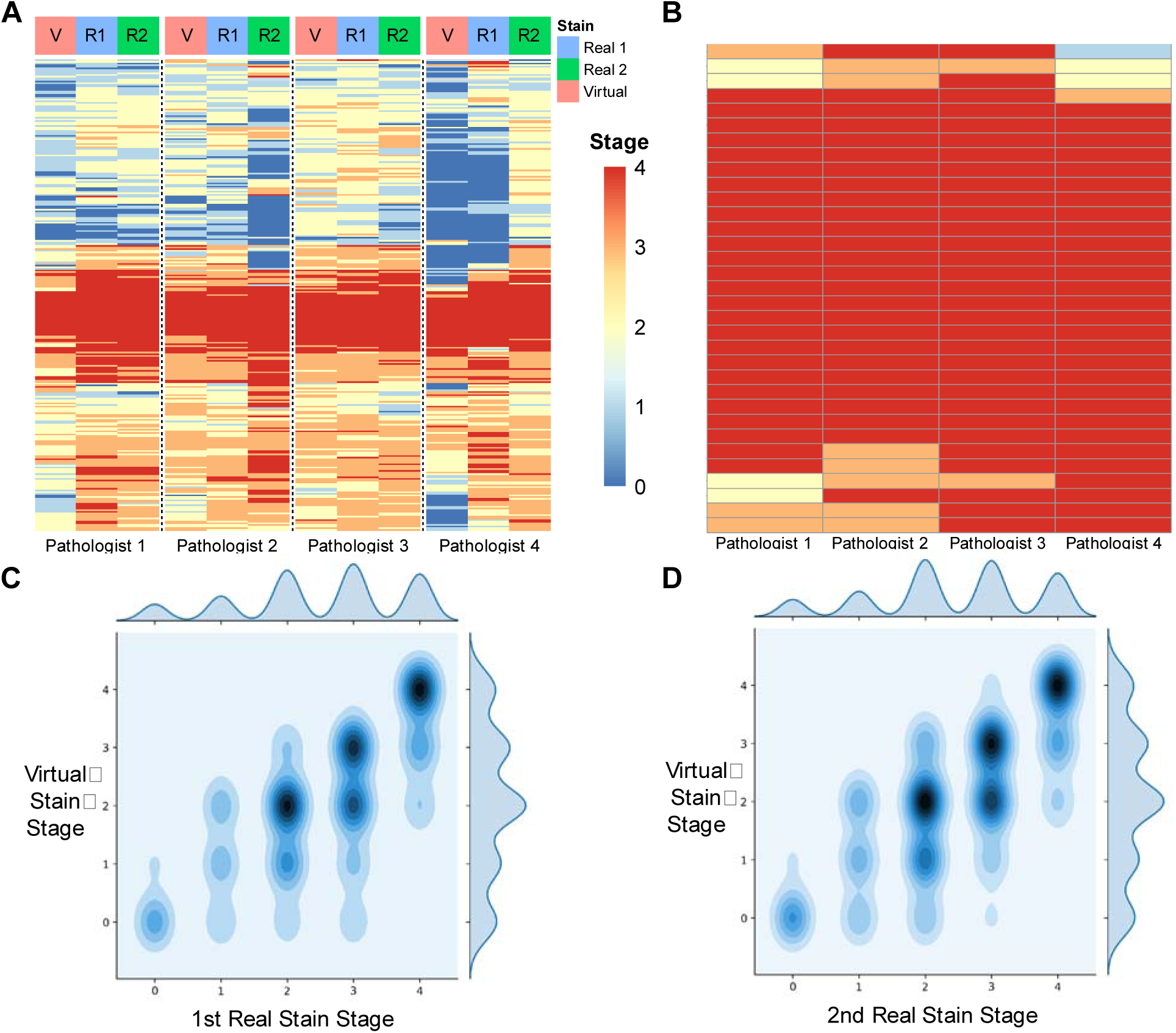
Concordance Plots Comparing Staging of Real to Virtual Stains; a) Heatmap of raw scores assigned by pathologists for real (blue column color for test one; green for test two) and virtual (red column color); each row represents a slide, each column represents a pathologist and whether they were staging real/virtual slide; set of each pathologist assessments (real one, real two, virtual) separated by column breaks; b) Similar heatmap (assessment of virtual stains) of agreement between pathologists but for slides deemed to have real advanced stage fibrosis (median real stage = 4.0; complete agreement in staging); c-d) 2-D Density plot of all stages (upper bound; F2-F3 to F3) assigned between real and virtual stains for the same slide; darker areas indicate higher concentration of stage combinations between real and virtual; off-diagonal elements have different staging between real and virtual images; c) virtual versus first assessment of real stains; d) virtual versus second assessment of real stains

Test-retest reliability (correlation) between assigned stages from subsequent inspections of the real stains for pathologists 1-4 were found to be 0.87 (95% CI: 0.84-0.90), 0.64 (95% CI: 0.59-0.70), 0.75 (95% CI: 0.71-0.80) and 0.55 (95% CI: 0.48-0.61) respectively. The average spearman correlation between stages of real stains across all pairs of pathologists was 0.69 from first set of real stages, while the average spearman correlation between pairs of pathologists was 0.70 from the second set of real stages. A complete set of inter-rater/intra-rater correlation in real staging and virtual staging can be found in the supplementary material (Supplementary Tables 5-6).

### Advanced Stage Fibrosis and Sensitivity Analysis

It is of paramount importance to detect advanced fibrosis with a high degree of sensitivity. Using slides where there was complete agreement between pathologists for advanced fibrosis (median *F* = 4.0; n=33 slides), staging of virtual slides recapitulated the advanced fibrosis (*F* = 4.0) staging 85% of the time (0.79, 0.79, 0.94, 0.88 for the pathologists respectively) (Figure 4b). Assessing agreement across all slides for both the presence (median *F* = 4) and absence (median *F* ≠ 4) of fibrosis yielded a Cohen’s Kappa of 0.76 (95% CI: 0.65-0.87), demonstrating strong concordance.

### Serological Analysis

While the results above demonstrate strong agreement in fibrosis stage between virtual and real trichrome staining, we hoped to correlate the assigned stages with serological markers of fibrosis and long-term effects of end stage liver disease to determine if the virtual stain correlates with disease severity. There were significant positive associations with markers of hepatocyte injury AST and ALT (p < 0.001) as well as hepatic synthetic function such as INR (p <0.001). Additionally, there was a strong association between virtual fibrosis stage and downstream effects of portal hypertension such as ascites (p=0.002). Lastly, real and virtual fibrosis stage were tested for correlation with non-invasive scoring systems MELD and Fib4. Interestingly, although we observed a trend toward correlation for both real and virtual stains with the MELD score, only the virtual stage met statistical significance criteria (p = 0.047). Both virtual and real stages were significantly associated with Fib4 score (p < 0.001). These results further support the validity of utilizing virtual trichrome stains for fibrosis staging (Supplementary Table 7).

Staging differences between virtual and real slides was also associated with key serological markers (Supplementary Table 7).

### Survival Analysis

Perhaps the most important indicator of whether the virtual stains can be meaningfully employed in the clinic is whether staging of these tissues are equally predictive of risk of progression to end stage liver disease.

We found that for both virtual and real stained tissue the NASH CRN fibrosis stages were equally predictive of occurrence of symptoms of end stage liver disease (ESLD) via Kaplan-Meier (KM) analysis (virtual trichrome log-rank p < 0.001, real trichrome log-rank p < 0.001, Figure 5). All KM estimates for treating staging as a continuous measure (median across pathologists, rounded to nearest integer) and dichotomous measure (median *F* ≥ 3.5) yielded p-values less than 0.0001 via log-rank statistic (Figure 6). The KM plots were consistent for dichotomized staging between real and virtual stains. Interestingly, the virtual trichrome fibrosis stage seemed to reflect the risk of progression to End Stage Liver Disease (ESLD) more accurately for the various fibrosis stages. Namely, the order of time to progression to ESLD for the real trichrome fibrosis stages is F4 < F2 < F1 < F3 < F0 while in the virtual trichrome fibrosis stages the order is F4 < F3 < F2 < F1 = F0. (Figure 6b).

**Figure 5:**
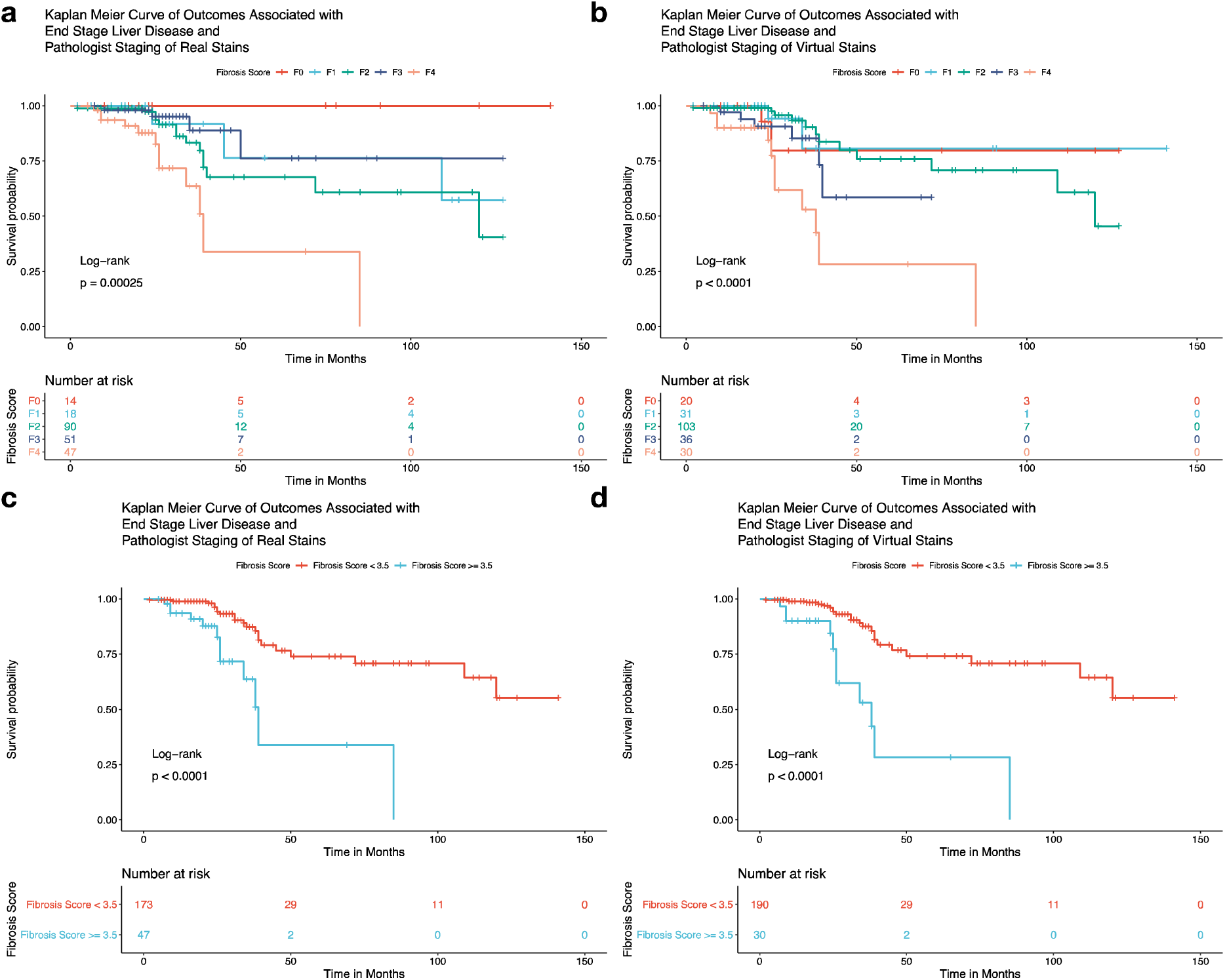
Kaplan-Meier Plots for the following conditions: (a,c) real stain staging; (b,d) virtual stain staging; (a,b) utilizing the median stage assigned by pathologists for that slide rounded to nearest integer; (c,d) dichotomizing staging by advanced fibrosis (*F* ≥ 3.5)

A Cox-Proportional Hazards (CoxPH) model for time-to-event outcomes adjusting for age and BMI was also applied which demonstrated an increased hazard of experiencing symptoms of end stage liver disease with increasing fibrosis stage in both real (Hazard Ratio HR = 2.06, CI 95% 1.36-3.12, P < 0.001) and virtually stained tissues (HR = 2.02, CI 95% 1.40-2.92, p < 0.001). The concordance statistic, which is a measure of fit of the survival curves to the data, was statistically equivalent for models fit on staging of real and virtual trichrome staging, irrespective of whether the staging was dichotomized. The computed hazard ratios were similar in effect and significance (Supplementary Tables 5-6). These trends, as elucidated via the KM and CoxPH analyses, were more pronounced when scores were dichotomized to advanced fibrosis (Fibrosis score greater than 3.5) or early fibrosis (score < 3.5). The aforementioned results suggest that virtual trichrome has the same ability to capture the potential for progression to ESLD as the real trichrome.

There were no statistically significant differences in dichotomized Fibrosis stage for both KM and CoxPH analyses of the real (KM p-value: 0.25, CoxPH p-value: 0.732) and virtual (KM p-value: 0.099, CoxPH p-value: 0.563) stain staging for the outcome of death.

We have summarized the results for concordance between pathologists staging scores for fibrosis. Overall, there was moderately strong reported agreement in scores from the real trichrome fibrosis stage to the stage derived from inspection of the virtual images. This indicates that, pending additional validation in the form of a prospective randomized controlled trial, virtual trichrome conversion can generate stains that can be readily deployed in clinical practice.

## Discussion

We investigated technologies that could potentially transition trichrome staining to a digital pathology test rather than a chemical stain, reducing the cost and personnel required to stage NASH biopsies. Our virtually stained tissue were visually consistent with real trichrome stained tissue as assessed by pathologists given the strong results from our Turing Tests, and our large-scale internal validation cohort quantitatively demonstrates strong correlation in staging between the real and virtual stains as assessed by pathologists. Virtually stained tissue fibrosis stages nearly surpassed our preliminary non-inferiority cutoff in concordance with two separate tests of real stage, had similar relationships with clinical / serological biomarkers, and were as predictive of progression to ESLD as their real trichrome stained counterparts. Our research presents a less labor and resource intensive solution to trichrome staining with nearly equivalent predictive value as traditional staining protocols. The ideal use of the technology as it currently stands is likely reflexive virtual trichrome staining for liver biopsies with real trichrome available by request (significantly reducing the volume of real trichrome stains performed), pending additional fine-tuning of the technology (Supplementary Figure 9).

Compared to other medical GAN studies, the strength in our design is the evaluation of clinical endpoints of the examined disease process. Our study, in addition to many others in the medical GAN literature, present “Turing Tests” that, while useful for evaluating the realism of generated images, have previously been utilized in the histopathology setting to assess small segments of tissue, which do not reflect whether the whole slide “looks real” ^39,66–68^. Though most of our sampled virtual subimages were indistinguishable from their real counterparts by a team of pathologists, we note that these measures of quality do not directly assess the viability of the generated slide for the potential to capture actionable clinical endpoints and place the pathologist in the position of evaluating a tiny subimage, rather than the entire slide. Accurately diagnosing and staging the presence of hepatic fibrosis is critical to the management of NASH. The presence of fibrosis, especially advanced fibrosis, has significant implications for medical management, frequency of disease assessment and prognosis. Additionally, accurate diagnostic information may serve as the impetus patients need to initiate lifestyle modifications and may indicate to the clinician the need for medication to manage their NASH risk factors ^69^. For these reasons it was extremely important for us to demonstrate, not only that our model could produce subjectively acceptable virtual trichrome subimages, but that it could create convincing WSI which were of sufficient quality to be accurately staged by pathologists. Given that this is an unproven technology that aims to replace or augment part of an already risky diagnostic procedure, we also thought it was necessary to take the additional steps of demonstrating that virtual trichrome fibrosis stage was strongly correlated with serological and clinical markers of liver injury and ESLD in a similar manner to real trichrome fibrosis stages. We were unable to make any substantive claims in our preliminary mortality-based survival analysis due to the relatively few patients who died during the study period ^2^. These factors merit further study of virtual staining in a cohort with greater representation of F4 stage tissue and sufficient mortality.

In general, the virtual trichrome images were deemed by the reviewers to be subjectively acceptable, confirmed by the use of Turing Tests. The most common comment recorded for the virtual stains was the presence of aberrant edge staining on needle core biopsies. This is likely the due to the inclusion of wedge biopsies in the training set. Because wedge tissue has significant areas of capsule, the model may have begun to associate any edge with trichrome positivity, even in the absence of capsular tissue ^70^. If this is the reason for the edge staining effect, then elimination of wedge biopsies with capsules in the final fine-tuning of our modeling approach may alleviate edge overstaining. Nonetheless, the raters in our study universally indicated that these areas of aberrant staining did not impede fibrosis scoring. Furthermore, quantitative results demonstrated that pathologists were able to recapitulate real late stage fibrosis using the virtual trichrome set.

### Limitations

There are several limitations to our work. The experiment was not completely blinded. Each pathologist was given a set of virtual trichromes first and then, two weeks later, staged the real set of trichromes. A final set of real stages were collected one to two months after staging the first real set. In order to minimize the chance of a pathologist recognizing a correspondent case between the real and virtual trichrome sets (“memorization”) we enforced a “washout” period between viewing the two sets of images and took the additional step of scrambling the case numbers and slide order. We acknowledge that this experimental design risks introducing bias, given that the raters might seek to judge the virtual images overly harshly or overly leniently and there is the small risk that the rater might remember their diagnosis from a virtual case while reviewing the analogous real case. However, given the limited time that raters were able to grant us, the goals we were seeking to accomplish (collecting subjective information about the virtual trichromes and rating fibrosis scores between real and virtual trichromes), and the limited number of pathologists with liver expertise available, we deemed these limitations acceptable. The results indicate that these potential biases may not have played a significant role, though we attempted to account for the aforementioned biases via our modeling approach regardless.

There was a high degree of inter-rater and intra-rater variability amongst assigned fibrosis stages from pathologists, regardless of whether the slide was real or virtual, as assessed from calculations of inter-rater correlation and test-retest reliability for real staging. The nature of the variability in scores was not random, as the final concordance modeling approach demonstrated the predisposition of some pathologists to statistically significantly assign a lower/higher virtual/real stage depending on their perception of the true stage of the patient. These behaviors may be related to the background of the clinician, such as age, time constraints, training, blinding, fatigue, personal preference, etc. That pathologists (and other visual experts) can render significantly different diagnoses for the same case (or even reviewing the same case at a later timepoint) has been well demonstrated in the literature ^71,72^ and represents a known limitation of qualitative diagnostics. For instance, the NASH CRN scale does not appear to be robust to changes in raters (which may be reflective of repeated assessments using a digital pathology medium over analog counterparts^71,73,74^; e.g. pathologists may have varying degrees of exposure with respect to this medium), which makes it a difficult to assess concordance to stages after use of a new staining technology, given the potential for high measurement error for the real stain stages.

In addition to this inter and intra rater variability, oftentimes the pathologist posted a range of stages (e.g. F2-F3) that represented their uncertainty in staging, which in turn made it difficult to model staging as an ordinal variable. These interval reports are reflective of the pathologist’s attempt to summarize an underlying continuous distribution of fibrosis stage. Many prior studies employing the NASH CRN system do not report such measures, though such approximations may yield meaningful information that should be taken into account when assessing the viability of a scale for clinical decision making. While we have begun to assess for these points of bias and uncertainty in our modeling approaches in this study (through utilization of a hierarchical Bayesian ordinal regression approach) the fitting procedures of such assessment models should likely be adapted to account for measurement uncertainty (i.e. considering an F2 and F3 at the same time rather than separately given an F2-3 report), though such procedures were beyond the study scope. Nonetheless, inclusion of hierarchical Bayesian modeling procedures are warranted to lend additional insight as to the assessment and validation of these technologies given the limitations of the evaluating scales and will be pursued in the future ^75,76^. Due to physician workload, we were only able to assess test-retest reliability, and concordance between the virtual stain stages and two successive sets of real stain stages; without a “ground-truth” to measure from and due to the risk of memorization of stage, we were thus unable to see whether their staging of real versus virtual tissue was within what was expected from intra-rater reliability. Prior studies have found merit in reporting a correlation based non-inferiority test^62^, which we had employed in our study to demonstrate preliminary agreement, but a more rigorous clinical readiness test of these technologies may be necessary to demonstrate equivalence and immediate usage of this technology ^77–80^.

### Future Opportunities

While these analyses were conducted on NASH cases, fibrosis can result from any chronic liver injury. Aside from NASH, chronic viral hepatitis, alcoholic steatohepatitis, autoimmune hepatitis, chronic biliary cirrhosis, primary sclerosing cholangitis and outflow obstruction are all relatively common causes of chronic liver injury. While fibrosis may be recognized equally well in these disease etiologies, the model needs additional testing on other manifestations of fibrosis unrelated to NASH which often have distinct scoring systems^81^. While virtual generation of a trichrome stain appears to be a viable solution for our medical center, we acknowledge that not all histological stains are amenable to virtual generation from H&E (eg. Ziehl Neelsen for acid-fast bacilli, iron stains). Furthermore, selection of routinely performed stains for the reporting of liver biopsies will vary between institutional labs (eg. Masson trichrome, PAS, Reticulin stains); we did not validate image-to-image translation techniques for inferring these stains but this presents future opportunity for other researchers.

While virtual staining may reduce labor costs and reagent usage, modern autostainers and laboratory automation efforts may also assist in this regard. Pathologists may be more likely to trust a real stain. However, the degree of laboratory automation, likewise computational workflow automation, remains variable between different institutions.

Furthermore, there are opportunities to improve on the modeling approach through registration and a paired analysis (e.g. Pix2Pix). Even when tissue is sectioned and captured to a slide perfectly (e.g. 5 microns between H&E and trichrome stain) there are significant alterations to the microstructure of the tissue (e.g. nuclei, cells and other small objects are caught in a different sectioning plane or are completely absent). Achieving perfect registration of an H&E and a trichrome stain for subsequent sections is therefore very challenging and rarely possible. Even if the H&E slide were destained and restained with trichrome, the chemical processing and handling involved (especially repeated cover slipping) introduces significant tissue distortions. Ultimately, factors related to the serial sectioning of tissue present significant disadvantages for training a stain conversion model using paired image information. We were not surprised when a preliminary evaluation of training a Pix2Pix model after registration of H&E and subsequent trichome sections demonstrated suboptimal results (principally significant structural laxity in the virtual images, so called ‘hallucination’) where the generated image was subjectively convincing but often bore no structural similarity to the source H&E. These difficulties may be alleviated through use of GAN techniques that allow imperfect registration, perhaps combining the best elements of the CycleGAN and Pix2Pix algorithms to build a more sophisticated model or incorporating new components into the loss term to enforce both structure preservation and capture overall similar tissue features.

There are obvious avenues to improving our current CycleGAN modeling approach including conducting a paired translation analysis, augmenting the number of capsule containing and capsule devoid training images (to address the edge staining artifact), and more examples capturing the center of large fibrotic nodules and portal areas (to address understaining seen in some large structures). In addition, the GAN generator was subjectively assessed (by comparing generated trichromes to their real counterpart) only every 5000 training iterations and there were often large differences between different training iterations. Saving more frequently and analyzing additional virtual WSI may therefore allow us to select an even better model for clinical use.

We have containerized our model and pre-processing pipeline into an opensource, deployable software package. These software elements can be readily incorporated into automated clinical workflows through the use of pipelining software such as Docker and Common Workflow Language (CWL) ^82,83^, and can be scaled to meet any clinical load with no additional coding (given the appropriate high-performance computing (HPC) resources). With additional fine-tuning the model presented herein could be readily deployed as a diagnostic aid. Pathologists could use this tool to flip rapidly between or compare images of H&E and trichrome slides (and other staining modalities), and if they are not satisfied with the staining quality of the virtual slide or find the interpretation ambiguous, can order a chemical stain. These steps can facilitate faster clinical decision making with less resources (both departmental and tissue). Future analyses may explore the prediction of fibrosis stage from an H&E, obviating the need for clinical inspection of a virtual stain ^84^. However, virtual staining is a technology that clinicians may trust more than a computer-generated risk score and therefore presents a viable decision aid technology.

## Conclusion

Our CycleGAN-based H&E to trichrome conversion model is interpretable and accurate and with appropriate validation could be incorporated into the clinical workflow as a diagnostic aid. The application of these techniques has the potential to save money, save time and improve patient outcomes by allowing for faster and more efficient fibrosis staging. In addition, methods like virtual trichrome staining offer a strong financial incentive for the adoption of digital pathology. We will investigate techniques to further refine this process while seeking implementation in the clinical workflow and prospective validation at DHMC.

## Abbreviations

ANN: Artificial Neural Network
ANOVA: One-Way Analysis of Variance
AUROC: Area Under the Receiver Operating Curve CNN – Convolutional Neural Network
CoxPH: Cox Proportional Hazards CWL – Common Workflow Language
DHMC: Dartmouth Hitchcock Medical Center ESLD – End Stage Liver Disease
GAN: Generative Adversarial Network H&E - Hematoxylin and Eosin
HE: Hazard Ratio
HPC: High Performance Computing HR – Hazards Ratio
IQR: Interquartile Range KM – Kaplan Meier
LMM: Linear Mixed Effects Model
NAFLD: Non-Alcoholic Fatty Liver Disease
NASH: Non-alcoholic Steatohepatitis ROC – Receiver Operating Curve
UNOS: United Network Organ Sharing WSI – Whole Slide Images

## Disclosures / Conflicts of Interest

The authors declare that they have no financial or non-financial conflicts of interest.

## Funding

This work was supported by:

- NIH grants R01CA216265, R01DE022772, and P20GM104416 to BCC
- Two Dartmouth College Neukom Institute for Computational Science CompX awards to BCC, LJV and JJL.
- JJL is supported through the Burroughs Wellcome Fund Big Data in the Life Sciences training grant at Dartmouth.
- Norris Cotton Cancer Center, DPLM Clinical Genomics and Advanced Technologies EDIT program

The funding bodies above did not have any role in the study design, data collection, analysis and interpretation, or writing of the manuscript.

## Authors’ Contributions

The conception and design of the study were contributed by JJL, BCC and LJV. Implementation, programming, data acquisition, and analyses were by JJL. Survival analysis, collection and analysis of serological markers and demographics were by NA. CB assisted with statistical consultation. AS, XL, ML, BR, and MA contributed to data collection and fibrosis staging. All authors contributed to writing and editing of the manuscript.

## Acknowledgements

Preliminary results for a small subset of samples featured in this study were presented at the COMP2CLINIC workshop at BIOSTEC 2020 ^23^. We would like to acknowledge Richard Zhan for his assistance in development of the openseadragon viewing framework for virtual trichrome stains. We would also like to acknowledge James O’Malley for statistical consultation.

## Notes

### Competing Interest Statement

The authors have declared no competing interest.

