## Supplementary Materials for "A Large-Scale Internal Validation Study of Unsupervised Virtual Trichrome Staining Technologies on Non-alcoholic Steatohepatitis Liver Biopsies"

**Supplementary Table 1: Demographic Characteristics; Organized by Median Fibrosis Staging of Real Cases**

|  | F0 | F1 | F2 | F3 | F4 | p |
| --- | --- | --- | --- | --- | --- | --- |
| n | 9 | 52 | 108 | 71 | 46 |  |
| Age (mean (SD)) | 48.56 (6.88) | 51.19 (11.56) | 52.94 (11.84) | 53.21 (14.67) | 55.80 (14.05) | 0.364 |
| Sex = Male (%) | 0 (0.0) | 12 (29.3) | 31 (34.8) | 24 (42.1) | 11 (27.5) | 0.422 |
| BMI (%) |  |  |  |  |  | 0.408 |
| Underweight (<20) | 0 (0.0) | 0 (0.0) | 2 (1.9) | 0 (0.0) | 0 (0.0) |  |
| Normal (20-25) | 1 (11.1) | 4 (7.7) | 6 (5.7) | 4 (5.6) | 6 (13.3) |  |
| Overweight (25-30) | 1 (11.1) | 9 (17.3) | 28 (26.7) | 13 (18.3) | 4 (8.9) |  |
| Obese (30-35) | 3 (33.3) | 17 (32.7) | 38 (36.2) | 22 (31.0) | 14 (31.1) |  |
| Severely Obese (>35) | 4 (44.4) | 22 (42.3) | 31 (29.5) | 32 (45.1) | 21 (46.7) |  |
| Impaired Fasting Glucose (%) | 4 (44.4) | 32 (61.5) | 59 (56.2) | 50 (70.4) | 36 (78.3) | 0.045 |
| Diagnosis of Diabetes Mellitus (%) | 2 (22.2) | 20 (38.5) | 40 (38.1) | 45 (63.4) | 27 (58.7) | 0.002 |
| Fib4 At Diagnosis (mean (SD)) | 2.61 (3.71) | 1.00 (0.61) | 1.80 (2.27) | 1.80 (1.29) | 2.90 (2.20) | <0.001 |
| Thrombocytopenia (%) |  |  |  |  |  | 0.022 |
| Normal | 5 (71.4) | 50 (98.0) | 87 (84.5) | 61 (87.1) | 31 (67.4) |  |
| Mild | 2 (28.6) | 1 (2.0) | 11 (10.7) | 7 (10.0) | 11 (23.9) |  |
| Moderate | 0 (0.0) | 0 (0.0) | 5 (4.9) | 2 (2.9) | 3 (6.5) |  |
| Severe | 0 (0.0) | 0 (0.0) | 0 (0.0) | 0 (0.0) | 1 (2.2) |  |
| INR Normal at Diagnosis (%) | 6 (100.0) | 19 (90.5) | 55 (100.0) | 43 (100.0) | 34 (91.9) | 0.053 |
| Albumin (%) |  |  |  |  |  | 0.093 |
| Normal (>3.5) | 7 (100.0) | 49 (100.0) | 94 (94.0) | 65 (92.9) | 39 (86.7) |  |
| Mild (2.8-3.5) | 0 (0.0) | 0 (0.0) | 2 (2.0) | 5 (7.1) | 3 (6.7) |  |
| Moderate (<2.8) | 0 (0.0) | 0 (0.0) | 4 (4.0) | 0 (0.0) | 3 (6.7) |  |
| Hyponatremic at Diagnosis (%) | 0 (0.0) | 2 (4.0) | 3 (2.9) | 3 (4.3) | 4 (9.3) | 0.504 |
| AST At Diagnosis(mean (SD)) | 131.86 (214.38) | 30.88 (19.86) | 51.77 (50.25) | 61.94 (47.70) | 64.11 (48.04) | <0.001 |
| ALT At Diagnosis (mean (SD)) | 177.29 (196.68) | 44.16 (32.95) | 64.38 (50.90) | 86.89 (75.37) | 63.24 (59.75) | <0.001 |
| Cr At Dx (mean (SD)) | 0.79 (0.19) | 0.87 (0.19) | 0.84 (0.23) | 0.90 (0.60) | 0.80 (0.25) | 0.586 |
| Billi At Dx (mean (SD)) | 1.87 (3.73) | 0.40 (0.16) | 0.62 (0.55) | 0.57 (0.75) | 0.86 (1.44) | 0.002 |
| Follow Up Time (mean (SD)) | 78.57 (46.69) | 34.72 (34.73) | 30.90 (23.65) | 31.81 (23.62) | 23.00 (14.67) | <0.001 |
| Fib4 at LCE (mean (SD)) | 1.02 (0.54) | 0.95 (0.38) | 1.48 (1.50) | 1.55 (1.03) | 2.82 (2.58) | <0.001 |
| Thrombocytopenia at LCE (%) |  |  |  |  |  | 0.041 |
| Normal | 6 (100.0) | 38 (97.4) | 72 (88.9) | 52 (91.2) | 21 (67.7) |  |
| Mild | 0 (0.0) | 1 (2.6) | 7 (8.6) | 3 (5.3) | 7 (22.6) |  |
| Moderate | 0 (0.0) | 0 (0.0) | 2 (2.5) | 1 (1.8) | 3 (9.7) |  |
| Severe | 0 (0.0) | 0 (0.0) | 0 (0.0) | 1 (1.8) | 0 (0.0) |  |
| INR Normal at LCE (%) | 0 (NaN) | 8 (100.0) | 37 (94.9) | 29 (90.6) | 27 (96.4) | NaN |
| Albumin at LCE (%) |  |  |  |  |  | 0.091 |
| Normal | 6 (100.0) | 39 (95.1) | 83 (96.5) | 56 (98.2) | 28 (87.5) |  |
| Mild | 0 (0.0) | 2 (4.9) | 2 (2.3) | 1 (1.8) | 1 (3.1) |  |
| Moderate | 0 (0.0) | 0 (0.0) | 1 (1.2) | 0 (0.0) | 3 (9.4) |  |
| Na at LCE(mean (SD)) | 141.40 (2.30) | 140.21 (2.54) | 140.29 (2.41) | 140.19 (2.61) | 139.03 (3.69) | 0.163 |
| Creatinine at LCE (mean (SD)) | 1.02 (0.25) | 0.80 (0.19) | 0.84 (0.27) | 1.02 (0.82) | 0.78 (0.21) | 0.067 |
| Bilirubin at LCE (mean (SD)) | 0.46 (0.42) | 0.48 (0.31) | 0.51 (0.31) | 0.57 (0.42) | 0.81 (0.97) | 0.032 |
| AST at LCE (mean (SD)) | 32.14 (11.77) | 24.21 (14.01) | 33.05 (30.17) | 37.79 (26.60) | 48.25 (32.16) | 0.004 |
| ALT at LCE (mean (SD)) | 49.14 (15.49) | 33.10 (27.81) | 36.53 (28.15) | 43.35 (35.81) | 46.06 (33.55) | 0.231 |
| HA1C at LCE (mean (SD)) | 5.92 (0.46) | 5.77 (1.10) | 6.11 (1.51) | 6.45 (1.73) | 6.41 (1.29) | 0.193 |
| Treated with Beta Blocker (%) | 0 (0.0) | 2 (4.0) | 4 (3.7) | 3 (4.2) | 3 (6.5) | 0.898 |
| Ascites (%) | 0 (0.0) | 0 (0.0) | 3 (2.8) | 3 (4.2) | 6 (13.0) | 0.019 |
| Evidence of Varices (%) | 0 (0.0) | 0 (0.0) | 7 (6.5) | 3 (4.2) | 6 (13.0) | 0.075 |
| Dialysis (%) | 0 (0.0) | 0 (0.0) | 1 (0.9) | 1 (1.4) | 1 (2.2) | 0.87 |
| Diagnosis of HCC 1 (%) | 0 (0.0) | 0 (0.0) | 0 (0.0) | 0 (0.0) | 2 (4.3) | 0.035 |
| Encephalopathy (%) | 0 (0.0) | 0 (0.0) | 1 (0.9) | 0 (0.0) | 5 (10.9) | <0.001 |
| Splenomegaly (%) | 1 (12.5) | 4 (8.2) | 6 (5.6) | 6 (8.5) | 8 (17.4) | 0.216 |
| Hospitalizations due to Liver Disease (%) | 0 (0.0) | 1 (2.0) | 7 (6.5) | 1 (1.4) | 5 (10.9) | 0.129 |
| Survival (% alive at LCE) | 9 (100.0) | 52 (100.0) | 102 (94.4) | 69 (97.2) | 42 (91.3) | 0.213 |

**Model Selection**

There are two major GAN translation techniques that may be utilized for stain translation depending on the type of data available, Pix2Pix or CycleGAN. Pix2Pix is able to translate one type of stain to another based on perfectly aligned paired images. Because of the one to one correspondence of pixels in the source and target domains, Pix2Pix is better able to retain the original image structure in the translation image.^1^ Analogously, for unpaired images, CycleGAN is utilized. Because the source and target images are unpaired and may show highly disparate scenes (e.g. a horse and a zebra in different poses), CycleGAN must compare the generated virtual/synthetic images to the real images and back translates to the original image to retain the original structure (see Supplementary Material section “Mathematical Description of CycleGAN”) ^2^. In our research, CycleGAN translated images tend to differ structurally from their source images more significantly than Pix2Pix translations, presumably due to the lack of perfectly paired source material.

We had to decide which of these two aforementioned image translation deep learning algorithms to utilize. The use of a Pix2Pix model would require paired images to be supplied and necessitate perfect alignment of the matching H&E and trichrome WSI. It proved too difficult to register WSI with enough fidelity for Pix2Pix, primarily as a result of tissue changes between adjacent sections ^3^. Thus, we opted to train a CycleGAN model due to imperfect registration ^3^ and different tissue features in adjacent tissue sections.

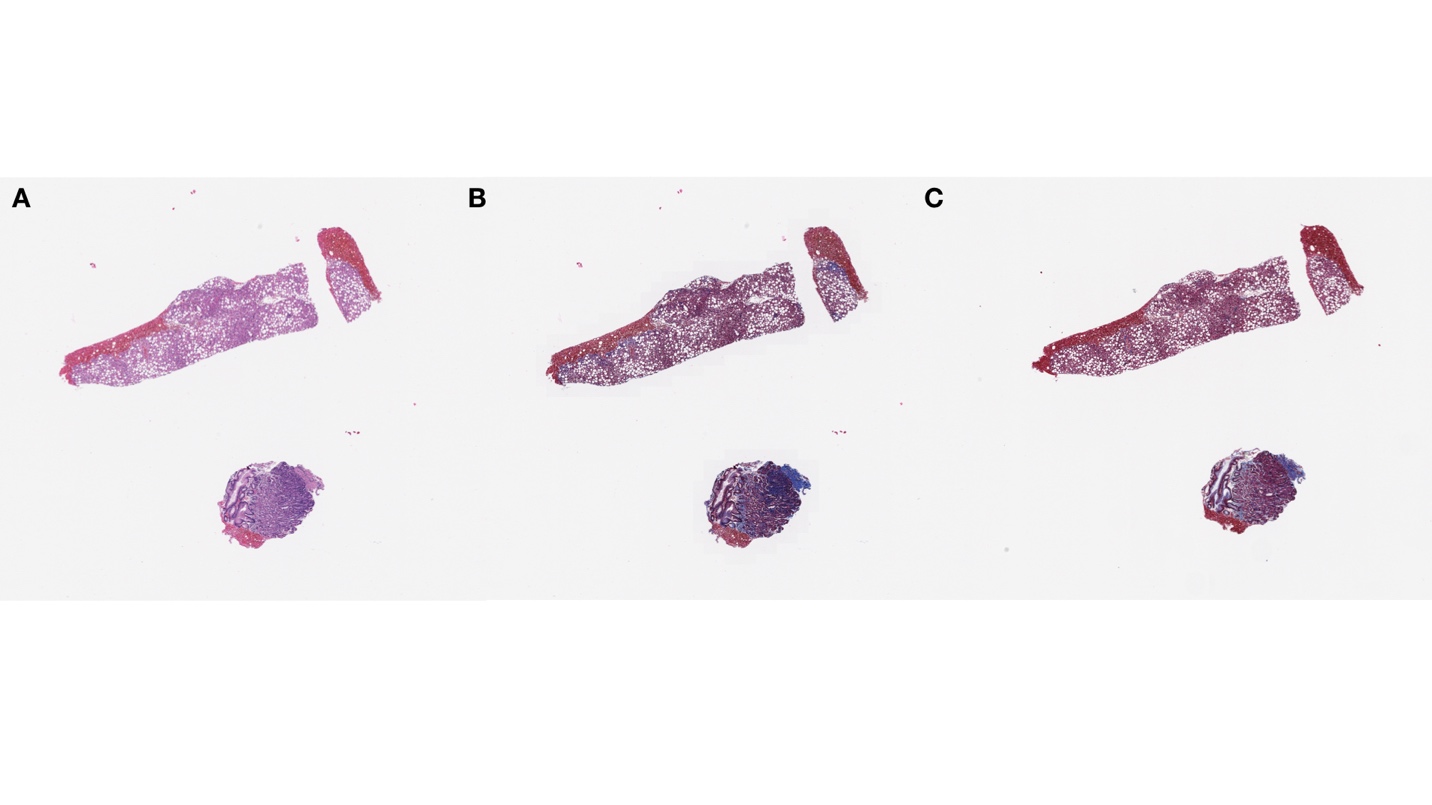

**Supplementary Figure 1: WSI for Slide ID 116 (F0):** a) H&E Staining; b) Virtual/Synthetic Trichrome Staining; c) Real Trichrome Stain from Adjacent Layer

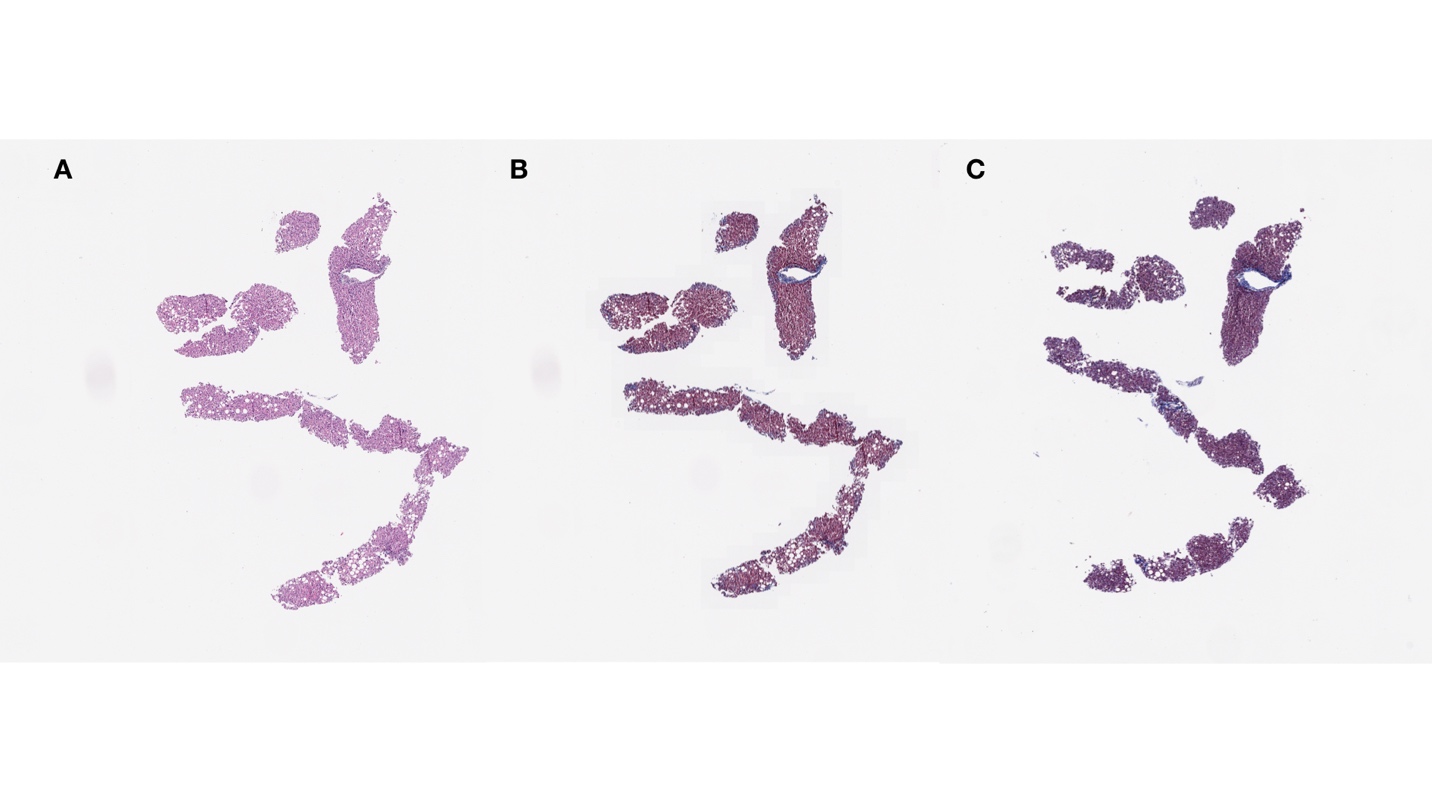

**Supplementary Figure 2: WSI for Slide ID 64 (F1):** a) H&E Staining; b) Virtual/Synthetic Trichrome Staining; c) Real Trichrome Stain from Adjacent Layer

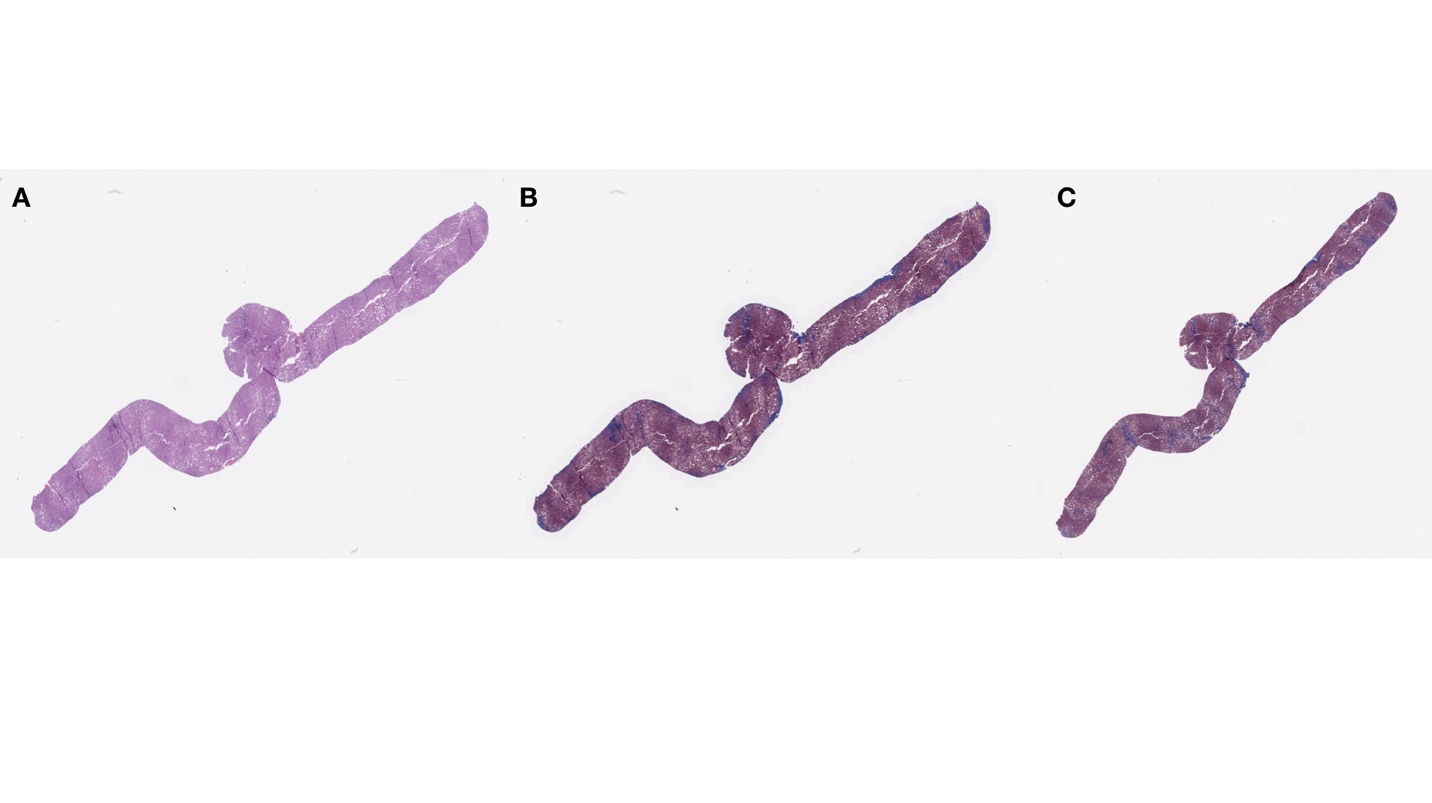

**Supplementary Figure 3: WSI for Slide ID 15 (F2):** a) H&E Staining; b) Virtual/Synthetic Trichrome Staining; c) Real Trichrome Stain from Adjacent Layer

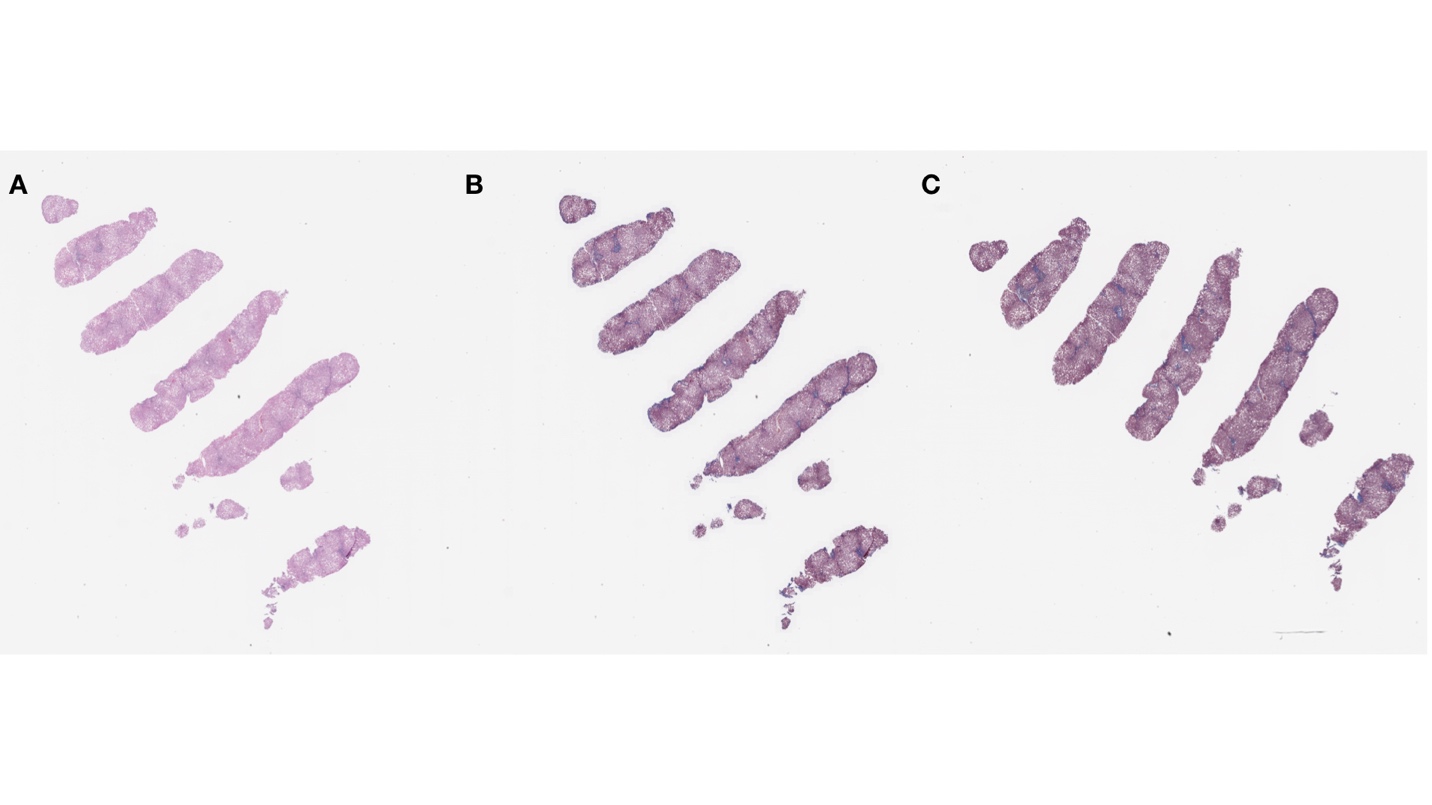

**Supplementary Figure 4: WSI for Slide ID 246 (F3):** a) H&E Staining; b) Virtual/Synthetic Trichrome Staining; c) Real Trichrome Stain from Adjacent Layer

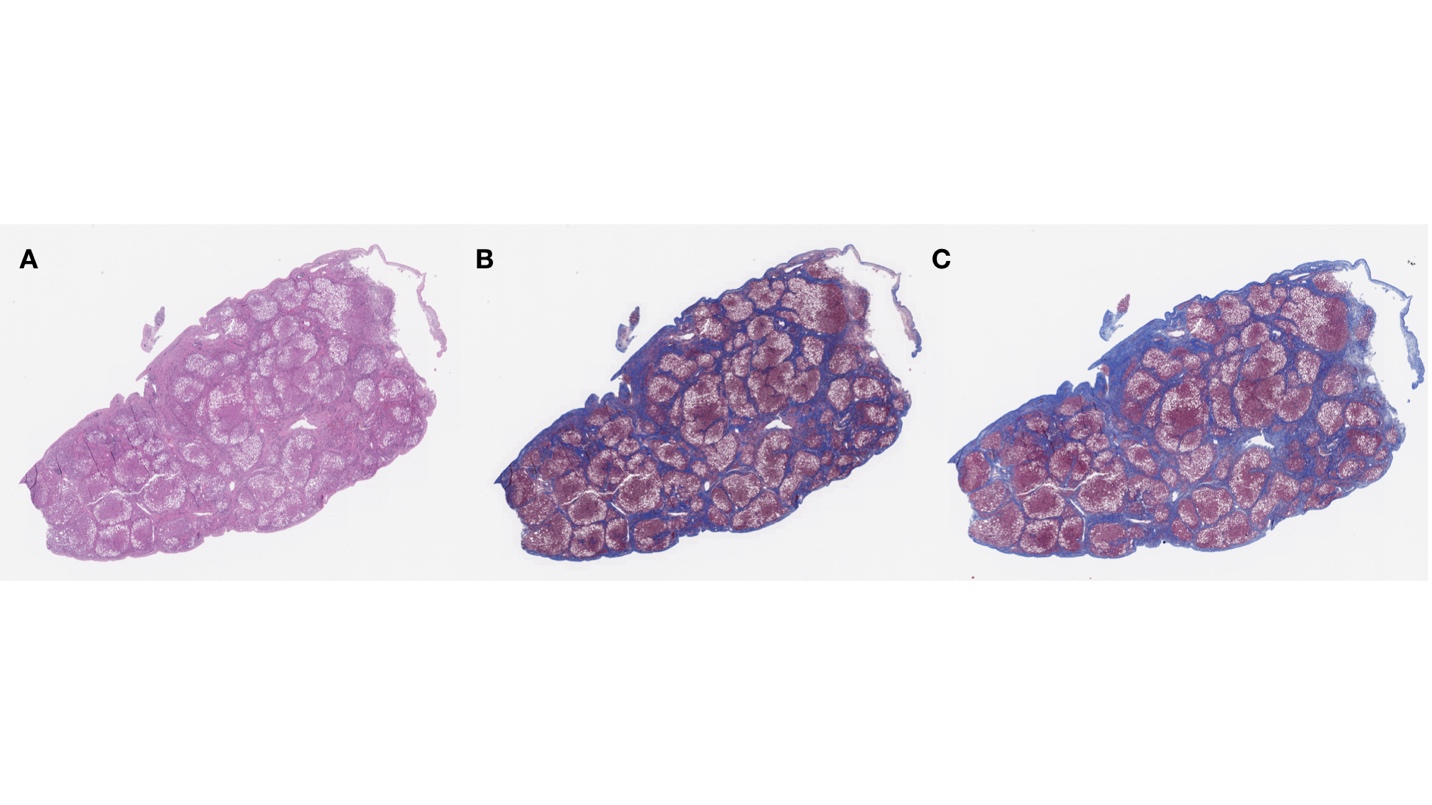

**Supplementary Figure 5: WSI for Slide ID 163 (F4):** a) H&E Staining; b) Virtual/Synthetic Trichrome Staining; c) Real Trichrome Stain from Adjacent Layer

**Supplementary Table 2:** Collection/Summary of pathologist comments on slides; a lot of the same comments mentioned for the real stains (e.g. fragmented), were also noted in the virtual stains, but there were additional artifacts introduced from virtual staining. In comparison to real stains (two slides from first set of inspection of real stains did not receive conclusive stage), all virtual stains were able to be staged.

Comments on Real Stains

Wedge/Needle biopsy

Fragmented biopsy (affected by biopsy procurement)

Rare Portal tract (suboptimal number of portal tracts for analysis)

Tangential section to capsule

Unable to Stage (n=2)

Additional Comments on Virtual Stains

Understaining

Understaining of capsules, with visible nodules

Overstaining of periphery regions

Blue hepatocytes

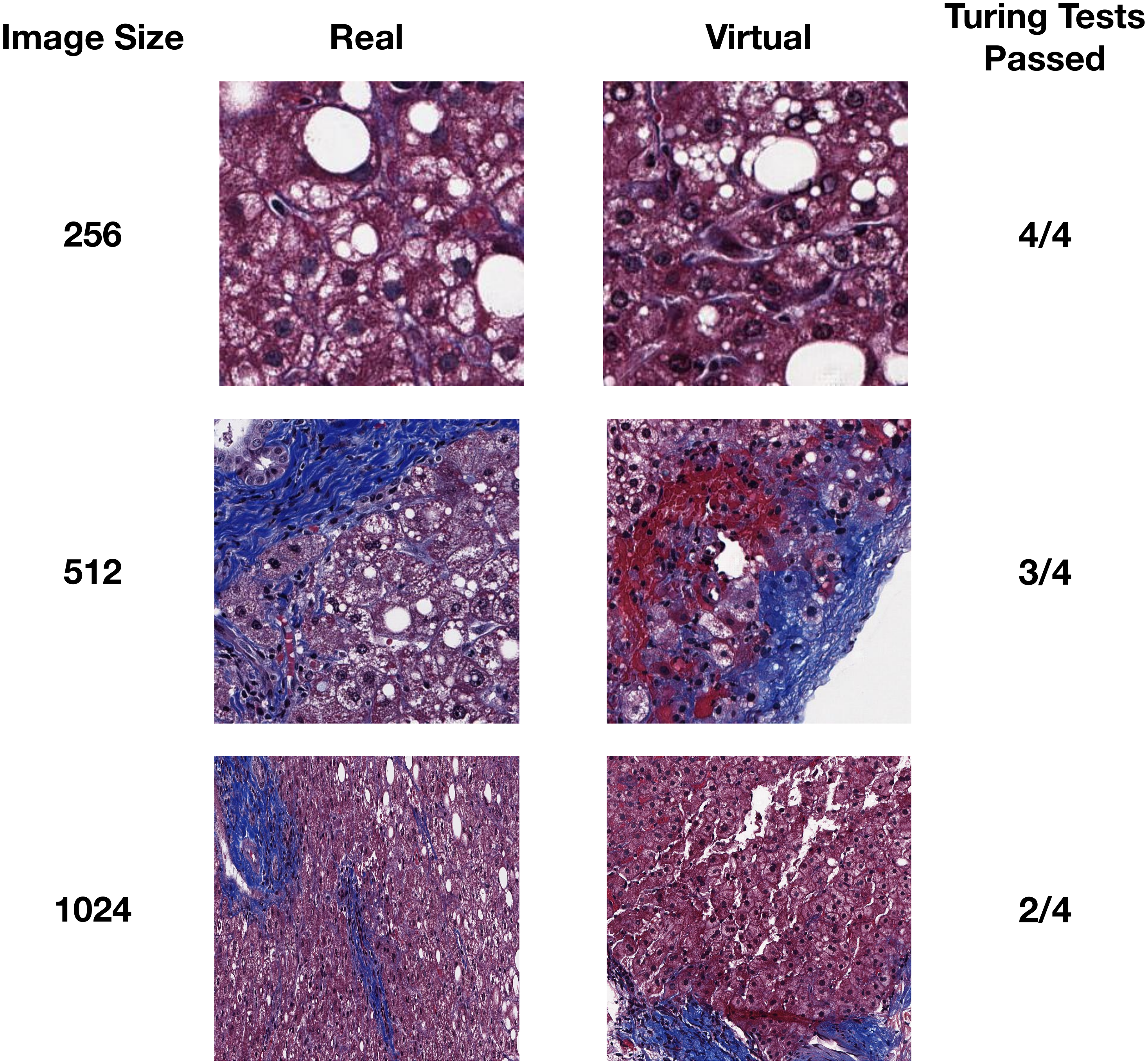

**Supplementary Figure 6:** Examples of images sampled from real and virtual stains for three image sizes (256 pixels, 512 pixels, 1024 pixels), correspondent to the twelve Turing Tests, the number of which passed has been enumerated per image size

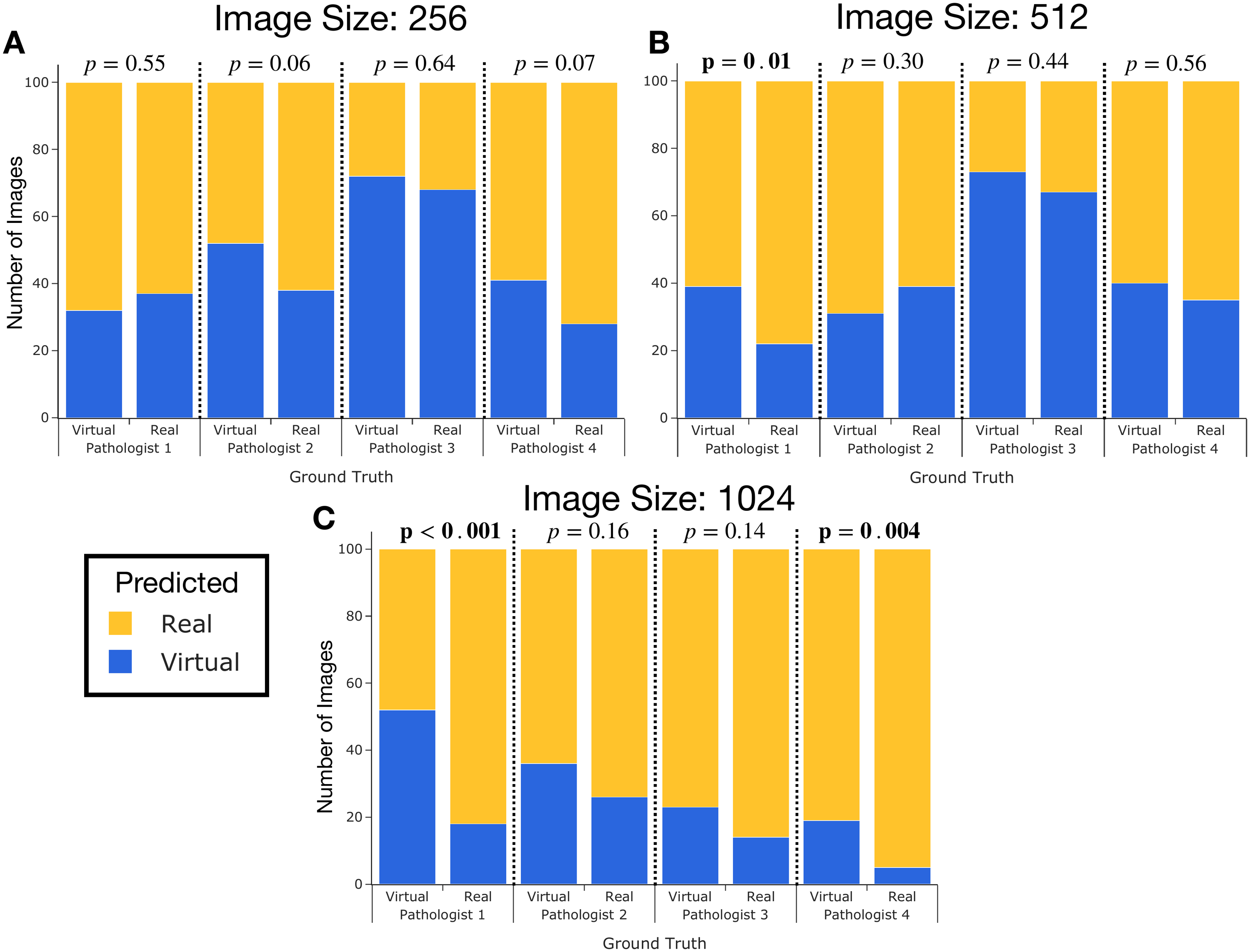

**Supplementary Figure 7:** Grouped stacked bar plots demonstrating results from twelve turing tests (n=2400 observations, 600 images assessed per pathologist; 4 pathologists; 100 real, 100 virtual per subimage size); yellow represents images assessed/predicted as real by pathologist; blue represents images assessed/predicted as virtual by pathologist; p-values greater than 0.05 demonstrate passing of turing test, where pathologists are unable to distinguish between real and virtual images; each bar represents for particular pathologist whether 100 images were actually real or virtual; if pathologists assessed similar amounts of images as virtual for images that are actually real and virtual, then pathologist was likely unable to tell the difference between real/virtual images (passed turing test); some were more likely to assign more images as virtual, regardless of whether image was real or virtual; where amount of blue significantly differs between two bars (actual real/virtual) for pathologist indicate instances where turing test did not pass, as indicated by the bolded p-value (Fischer’s exact test); these procedures were done for images of size: a) 256; b) 512; c) 1024

**Supplementary Table 3:** Tabulation of Turing Test results, indicating same data from bar plots in Supplementary Figure 7; two by two contingency matrices were calculated for each image size (x3) and for each pathologist (x4), yielding twelve contingency tables of number of images predicted by pathologist to be real or virtual versus the actual number of images that were real or virtual; columns denote whether image was assessed to be real or virtual by pathologist; rows denote whether image was actually real or virtual; numbers were bolded if they came from a failed turing test

|  | | Predicted by Pathologist | | | | | | | | | | |
| --- | --- | --- | --- | --- | --- | --- | --- | --- | --- | --- | --- | --- |
|  | | **Pathologist 1** | |  | **Pathologist 2** | |  | **Pathologist 3** | |  | **Pathologist 4** | |
| Image Size (pixels) | **Ground Truth**  $\downarrow$ | **Virtual** | **Real** |  | **Virtual** | **Real** |  | **Virtual** | **Real** |  | **Virtual** | **Real** |
| 256 | **Virtual** | 32 | 68 |  | 52 | 48 |  | 72 | 28 |  | 41 | 59 |
|  | **Real** | 37 | 63 |  | 38 | 62 |  | 68 | 32 |  | 28 | 72 |
| 512 | **Virtual** | **39** | **61** |  | 31 | 69 |  | 73 | 27 |  | 40 | 60 |
|  | **Real** | **22** | **78** |  | 39 | 61 |  | 67 | 33 |  | 35 | 65 |
| 1024 | **Virtual** | **52** | **48** |  | 36 | 64 |  | 23 | 77 |  | **19** | **81** |
|  | **Real** | **18** | **82** |  | 26 | 74 |  | 14 | 86 |  | **5** | **95** |

**Challenges with Assessment via NASH CRN System and Justification for Statistical Modeling Approach**

Fibrosis stage, under the NASH CRN system, is an ordinal measurement, which presents various challenges when assessing concordance. While the scale attempts to approximate an underlying continuous measure of fibrosis, transitions between an F0 and an F1 may be entirely different from a transition between an F3 and F4, warranting a modeling approach that accounts for stage as an ordered categorical variable. Furthermore, the stage is bound between F0 and F4, which means that interrater variability in scoring the real slides may peak in intermediate stages, where the degree of fibrosis is more ambiguous, and subsequently more prone to measurement error and pathologist bias. The NASH CRN scale has demonstrated a limited interrater concordance for the task of fibrosis staging^4^, which is a measure generally associated with measurement error ^5^; this places an upper limit on what demonstrates reasonable concordance. As compared to other prior non-inferiority tests which have compared diagnosis between microscopy and WSI modalities, the non-inferiority test is conducted entirely on a digital medium a domain in which the expert pathologists have less experience compared to traditional microscopy. Finally, the expert pathologists reported a significant number of interval measurements (eg. F0-F1, F2-F3; 521 individual observations correspondent to 67% of the slides featured interval measurements) for real and virtual stages, which convey their uncertainty in outcome and their attempt at summarizing/approximating an underlying continuous distribution of degree of fibrosis spread that remains unmeasured as per the ordinality of the NASH CRN system. Such interval measurements have been unreported in prior characterizations of the NASH CRN system, yet convey meaningful information and similar interval scores are very commonly seen in other ordinal systems in pathology. A method to fully characterize this uncertainty by utilizing both measurements (eg. F2 and F3) simultaneously to capture more meaningful estimate of fibrosis progression is outside of the study scope.

**Treatment of Pathologist Staging Uncertainty for Study Evaluation**

Our group was tasked with establishing modeling procedures for ordinal variables that takes into account the interval measurements discussed in the previous section (eg. F2-F3). Had we forced pathologists to select between the two listed stages (eg. selecting F2 instead of F3), we would be discarding meaningful information from the modeling approach that more closely approximates their true assessment of fibrosis progression. There do not yet exist statistical modeling procedures that incorporate both measurements into an assessment model for NASH CRN staging or the system’s application for medical artificial intelligence technologies, which would provide a more accurate assessment of concordance of virtual to real stages. Moreover, the discussion of such estimation procedures would detract from the results of the study. Instead, we opted to consider separately the situations in which the lower bound (eg. F2) was considered, and in which the upper bound was considered (eg. F3) across all slides (given the report F2-3). This was done for the correlation tests in the text between virtual and real staging. The results from the correlation between real-virtual analysis (see “Statistical Methods and Results”) may be coarsely interpreted by observing the minimum and maximum correlation achieved across all statistical tests, though in the main text we considered results from the test that considers the upper bound of uncertain measurements, which we believe is more sensitive to advanced stage fibrosis.

We felt that considering the upper and lower bounds of fibrosis stages was not warranted for the rest of the study because the estimates when considered across all slides should yield similar results. As such, we considered the upper bound of uncertain fibrosis stages for the tests of agreement for advanced stage fibrosis, serological markers, and survival models due to heightened sensitivity for advanced stage fibrosis (eg. F3-F4 becomes an F4) ^6^.

**Statistical Methods and Results**

We assessed bias-adjusted correlation between real and virtual stage using an ordinal regression framework.

The dependent variable, virtual staging, is modeled using the cumulative link model^7,8^, which considers the cumulative distribution of the response. Let the response $Y_{i}$ denote virtual fibrosis stage. The probability that fibrosis stage falls up to stage $k\in\{0, 1, 2, 3, 4\}$ is $\alpha_{ik}=P\left( Y_{i}\leq k \right)=\sum_{j=0}^{j=k} P\left( Y_{i}=j \right)$. The cumulative link model assumes that stage $k$ is an actualization or indicator of an underlying/latent continuous variable that is representative of true fibrosis progression, $Y_{i}^{*}$, modeled as a continuous variable and following the following linear model: $Y_{i}^{*}=x_{i}\beta+\epsilon_{i}$, where $x_{i}$ represents the other covariates and independent variable. The five possible stages may be represented as five points on this continuous scale after being transformed via a link function $g$: $g\left( \alpha_{ik} \right)=\theta_{k}+x_{i}\beta$, where $\theta_{k}$ ($k\in\{0, 1, 2, 3\}$) represents a baseline or “cutpoint” of this continuous distribution; observed stages fall between these cutpoints. When making predictions, if $Y_{i}^{*}$ is less than $\theta_{0}$, the stage 0 is predicted, and if $\theta_{0}<Y_{i}^{*}<\theta_{1}$, the stage 1 is predicted, etc. For the transformation that maps an ordinal variable to its underlying continuous distribution, the proportional odds (logistic regression extension; $g=logit$) or probit model ($g=\Phi^{-1}$) link functions may be considered.

The independent variable, real staging (utilized as a predictor to assess bias given perception of true stage because of its possible interaction with a given pathologist)^9^, is modeled as a monotonic effect to take into account ordinality^10^, because utilizing the variable as continuous does not contain information about the distance between adjacent categories. Monotonic effects attempt to more accurately capture information about the distance between adjacent categories by representing stage as a sum of increasing real numbers. If we consider $X_{i}$ to be real stage, where $x_{i}\in\{0,1,2,3,4\}$, and if $x_{i}$ was of stage k, $x_{i}\to mo\left( x_{i},\vec{\zeta} \right)=k\sum_{j=0}^{j=k} \zeta_{j}$, where $\sum_{j=0}^{j=4} \zeta_{j}$=1 and $\zeta_{j}\in\left[ 0,1 \right]$. The simplex $\vec{\zeta}$ is a vector that represents the distances between ordinal categories. In other words, an ordinal predictor can be represented in such a way that the ordinality is preserved, while learning and respecting the distances between the categories (eg. distance between an F3-F4 is meaningfully represented, parameterized by $\zeta_{3}$).

After inclusion of hierarchical modelling framework that accounts for repeated measures for a particular case, which induces correlated observations between pathologists for a given case, combining the cumulative link model which describes an ordinal outcome (virtual stage) and the monotonic effect which describes an ordinal predictor (real stage), and including interaction effects, we arrive at the specification for our statistical model:

$$virtual stage\sim real stage*pathologist+\left( 1+test | case \right)$$

Where *virtual stage* is $Y_{i}$, the ordinal response via the cumulative link model, *real stage* is $X_{i}$, the monotonic effect, *pathologist*, $\vec{Z_{i}}$ is an indicator vector for the chosen *pathologist*, and *test*, $t_{i}$, is an variable $t_{i}\in\{1,2\}$ indicating whether the real stage came from the first or second test for real stages.

We utilized a bayesian modeling procedure to fit the described model, using Markov Chain Monte Carlo (MCMC) procedures which iteratively samples the posterior distribution of parameters until the joint posterior converges on a stationary solution; sampling may be conditional on the posterior distribution of the other parameters (Gibb’s sampling) or trajectory-based (Hamiltonian Monte Carlo)^11^. Bayesian modeling procedures are often advantageous as compared to their frequentist counterparts because estimates are often more precise, less biased, computations are easier to implement and require fewer assumptions about the shape of the distribution of the estimated parameters. We utilized non-informative priors for each of the model parameters, selecting default priors supplied by the *brms* R software package^12^. Markov chains were sampled using a warmup period of 2000 iterations and then sampled for an additional 1000 iterations using the No-U-Turn-Sampler (NUTS) via the *brms* package^13^. Estimates of bias-adjusted real stage to correlate with virtual stage were obtained through sampling of the posterior distribution of parameter estimates and then applying the sampled parameters to the cumulative link model to yield probabilities for obtaining a particular stage. This predictive posterior distribution was averaged and the stage with the highest probability was selected as the bias-adjusted real stage. These bias-adjusted real-stage measures were correlated to the virtual stage via a non-parametric bootstrap of spearman’s correlation coefficient.

The R code utilized to fit the hierarchical bayesian ordinal model can be found below ^5,12^:

library(brms)

brm(virtual~mo(real)*rater+(1+test|case), data=design.matrix, family=cumulative(),cores=4,warmup = 2000, iter = 3000)

We have included 95% credible intervals (95% of the posterior distribution of the parameter lies in this interval) for many of the important parameters of the provided model in the following figure and have included the final estimates in the subsequent supplemental table. We fit four separate models corresponding to whether we restricted inter-rater variability (yes/no) and whether we utilized the upper bound of ambiguous staging (eg. F3 for F2-F3 report; yes/no).

**Supplementary Figure 8:** Posterior estimates of important main effects from Hierarchical Bayesian ordinal regression models under the following specifications: a, c) utilized all samples from inspection of virtual stains and test and re-inspection of real stains; d, b) incorporated samples/slides with low inter-rater variability as defined by the $IQR\leq3$; c,d) utilized upper bound in ambiguous staging assignments (eg. F3-F4 changed to F4) for real and virtual stages; a,b) utilized lower bound in ambiguous staging assignments (eg. F3-F4 changed to F3); dot represents mean of posterior distribution of parameter; thick line represents 50% credible interval; thin line represents 95% credible interval; *Cutpoints* represent various baseline measurements of latent variable of continuous fibrosis progression that are used to determine the probability of observing a certain stage (eg. if the latent variable is observed to be between *Cutpoint 1* and *Cutpoint 2*, it is likely a stage 1, etc..); the parameter corresponding to monotonic real stage representing the direction and strength of the correspondence between real stage and virtual, individual parameters corresponding to monotonic simplices $\zeta$ that represent the differences between stages were included in the model but not included in this visualization

**Supplementary Table 4:** Bootstrapped bias-adjusted spearman correlation estimates between real stage and virtual stage; estimated under the following scenarios (reflected by model estimates from Supplementary Figure 8): a, c) utilized all samples from inspection of virtual stains and test and re-inspection of real stains; d, b) incorporated samples/slides with low inter-rater variability as defined by the $IQR\leq3$; c,d) utilized upper bound in ambiguous staging assignments (eg. F3-F4 changed to F4) for real and virtual stages; a,b) utilized lower bound in ambiguous staging assignments (eg. F3-F4 changed to F3); estimates c-d were reported in the main text due to sensitivity to advanced stage fibrosis

| **Supplementary Figure 8 Letter** | **Restrict Inter-rater Variability?** | **Upper Bound of Ambiguous Assigned Stage** | **Number Observations** | **Spearman Correlation** | **2.5% Quantile** | **97.5% Quantile** |
| --- | --- | --- | --- | --- | --- | --- |
| A | No | No | 2291 | 0.80 | 0.78 | 0.82 |
| B | Yes | No | 1106 | 0.81 | 0.78 | 0.83 |
| C (in main text) | No | Yes | 2291 | 0.81 | 0.79 | 0.83 |
| D (in main text) | Yes | Yes | 1107 | 0.86 | 0.84 | 0.88 |

**Supplementary Table 5:** Spearman correlation matrix representing inter-rater correlation in staging real stains and test-retest reliability between subsequent inspections of real stains; correlations reported in upper diagonal (blue) represent correlation between pairs of pathologists for staining reals during the first pass inspection after the first washout period; correlations reported in lower diagonal (orange) represent correlation between pairs of pathologists for staining reals during the second pass inspection after the second washout period; correlations on the diagonal represent intra-rater correlation in staging between first and second assessment of real stains

|  | | **Test 1** | | | |
| --- | --- | --- | --- | --- | --- |
|  |  | **Pathologist 1** | **Pathologist 2** | **Pathologist 3** | **Pathologist 4** |
| **Test 2** | **Pathologist 1** | 0.87 | 0.82 | 0.74 | 0.64 |
|  | **Pathologist 2** | 0.64 | 0.64 | 0.75 | 0.64 |
|  | **Pathologist 3** | 0.82 | 0.59 | 0.75 | 0.53 |
|  | **Pathologist 4** | 0.80 | 0.60 | 0.78 | 0.55 |

**Supplementary Table 6**: Spearman correlation matrix representing inter-rater correlation in staging virtual stains

|  | Pathologist 1 | Pathologist 2 | Pathologist 3 | Pathologist 4 |
| --- | --- | --- | --- | --- |
| Pathologist 1 | -- | -- | -- | -- |
| Pathologist 2 | 0.74 | -- | -- | -- |
| Pathologist 3 | 0.68 | 0.66 | -- | -- |
| Pathologist 4 | 0.61 | 0.65 | 0.57 | -- |

**Supplementary Table 7:** Spearman Correlation of Various Staining Measures with Clinical Biomarkers; a) Median Real Trichrome Stage; b) Median Virtual Trichrome Stage; c) Difference between Median Virtual Stage and Median Real Stage; d) Interquartile Range (IQR) between Pathologists for Real Stage; Interquartile Range (IQR) between Pathologists for Virtual Stage

|  | **Median Real Trichrome Stage** | | **Median Virtual Trichrome Stage** | | **Median Residual Stage** | | **IQR in Real Staging** | | **IQR in Virtual Staging** | |
| --- | --- | --- | --- | --- | --- | --- | --- | --- | --- | --- |
|  | **ρ** | **p-value** | **ρ** | **p-value** | **ρ** | **p-value** | **ρ** | **p-value** | **ρ** | **p-value** |
| **Fib4** | 0.37 | <0.01 | 0.43 | <0.01 | 0.12 | 0.06 | -0.27 | <0.01 | -0.31 | <0.01 |
| **INR** | 0.25 | <0.01 | 0.27 | <0.01 | 0.11 | 0.17 | -0.13 | 0.11 | -0.14 | 0.08 |
| **ALB** | -0.09 | 0.14 | -0.03 | 0.57 | 0.03 | 0.63 | -0.03 | 0.58 | 0.09 | 0.12 |
| **AST** | 0.28 | <0.01 | 0.33 | <0.01 | 0.06 | 0.34 | -0.28 | <0.01 | -0.24 | <0.01 |
| **ALT** | 0.08 | 0.16 | 0.17 | <0.01 | 0.09 | 0.12 | -0.19 | <0.01 | -0.09 | 0.12 |
| **AST:ALT Ratio** | 0.26 | <0.01 | 0.20 | <0.01 | -0.07 | 0.22 | -0.06 | 0.32 | -0.17 | 0.01 |
| **Sodium** | 0.03 | 0.60 | -0.08 | 0.21 | -0.14 | 0.02 | 0.00 | 0.96 | 0.06 | 0.31 |
| **Hospitalization** | 0.07 | 0.24 | 0.09 | 0.12 | 0.05 | 0.43 | -0.13 | 0.03 | -0.15 | 0.01 |
| **Ascites** | 0.19 | <0.01 | 0.17 | <0.01 | 0.06 | 0.28 | -0.17 | <0.01 | -0.11 | 0.07 |
| **Alive** | -0.09 | 0.13 | -0.07 | 0.21 | 0.03 | 0.59 | 0.05 | 0.44 | 0.08 | 0.16 |
| **Variceal Bleed** | 0.03 | 0.61 | 0.06 | 0.30 | 0.03 | 0.61 | -0.07 | 0.21 | -0.05 | 0.42 |
| **BMI** | 0.07 | 0.27 | -0.06 | 0.35 | -0.17 | <0.001 | 0.06 | 0.34 | -0.01 | 0.85 |
| **Age** | 0.14 | 0.02 | 0.18 | <0.01 | 0.09 | 0.12 | -0.07 | 0.24 | -0.11 | 0.05 |
| **iMeld** | 0.16 | 0.05 | 0.21 | 0.01 | 0.16 | 0.04 | -0.16 | 0.04 | -0.05 | 0.56 |
| **Meld-Na** | 0.20 | 0.01 | 0.28 | <0.01 | 0.17 | 0.03 | -0.14 | 0.08 | -0.09 | 0.25 |

**Supplementary Table 8:** Model Fit Results from Cox-Proportional Hazards Models (adjusted for age/BMI) for median assigned stage

|  | **Real Trichrome** | | | **Virtual Trichrome** | | |
| --- | --- | --- | --- | --- | --- | --- |
| *Predictors* | *Hazard Ratio* | *CI* | *p* | *Hazard Ratio* | *CI* | *p* |
| Age | 1.02 | 0.99 – 1.05 | 0.133 | 1.02 | 0.99 – 1.04 | 0.298 |
| BMI | 0.91 | 0.86 – 0.98 | **0.007** | 0.93 | 0.87 – 0.99 | **0.** **017** |
| Median Stage | 2.06 | 1.36 – 3.12 | **<0.001** | 2.02 | 1.40 – 2.92 | **<0.001** |
| Observations | 220 | | | 220 | | |
| R^2^ Nagelkerke | 0.488 | | | 0.517 | | |
| Concordance (±SE) | 0.7324±0.0467 | | | 0.7365±0.0513 | | |

**Supplementary Table 9:** Model Fit Results from Cox-Proportional Hazards Models (adjusted for age/BMI), dichotomizing median assigned staging by advanced fibrosis ($F\geq3.5$)

|  | **Real Trichrome** | | | **Virtual Trichrome** | | |
| --- | --- | --- | --- | --- | --- | --- |
| *Predictors* | *Hazard Ratio* | *CI* | *p* | *Hazard Ratio* | *CI* | *p* |
| Age | 1.02 | 0.99 – 1.05 | 0.105 | 1.02 | 0.99 – 1.05 | 0.180 |
| BMI | 0.92 | 0.86 – 0.98 | **0.013** | 0.93 | 0.86 – 0.98 | **0.015** |
| Median F $\geq$3.5 | 5.02 | 2.28 – 11.05 | **<0.001** | 5.36 | 2.48 – 11.58 | **<0.001** |
| Observations | 220 | | | 220 | | |
| R^2^ Nagelkerke | 0.513 | | | 0.524 | | |
| Concordance (±SE) | 0.7541±0.04617 | | | 0.7502±0.0534 | | |

**Mathematical Description of CycleGAN**

Generative adversarial networks (GANs) generate realistic images, $X$, from a stochastic noise vector $Z$ via the neural network mapping: $g: Z\to X$. The process of training the generator, $G$, to produce a realistic image, $\hat{X}=G\left( Z \right)$, that matches the distributional properties of real images $X$ involves the use of a discriminator, $D$, which decides whether the produced and real images are real or virtual/synthetic. To do this, the discriminator estimates the low dimensional probability distribution of $X$ and $\hat{X}$. During optimization of the objective function for the model, the generator seeks for the density of $\hat{X}$ to match $X$, while the discriminator seeks to tease apart these two distributions. The model objective is the divergence between these two distributions may be encoded via the cross-entropy loss or alternatively the symmetrized Kullback-Leibler divergence (Jensen–Shannon Divergence), reformulated as such:

$$L_{GAN}\left( G,D \right)=E_{x\sim p_{r}\left( x \right)}\left[ log\left( D\left( x \right) \right) \right]+E_{x\sim p_{g}\left( x \right)}\left[ log\left( 1-D\left( G\left( z \right) \right) \right) \right]$$

The real distribution of data is $p_{r}\left( x \right)$ while the generated distribution is $p_{g}\left( x \right)$. The parameters of the generator are estimated through the alternating updates to the generator and discriminator via the *minimax* game:

$${G^{*}=argmin_{G}\mathrm{ma}x_{D}L}_{GAN}\left( G,D \right)$$

The minimization of the loss with respect to the generator parameters via gradient descent only operates on the right-hand term of the objective, which updates the parameters of the generator such that the discriminator estimates a higher probability for a generated sample of being real. The maximization of the loss with respect to the discriminator parameters attempts to maximize the probability of a real sample being real on the left-hand of the equation while minimizing the probability of a generated sample being real on the right-hand of the equation. The game that is played between the generator and discriminator may cause the loss to oscillate during training and is unstable, warranting further modifications to the objective function (eg. Wasserstein distance and gradient penalties) and demands close monitoring and visual inspection to ensure the fidelity of the samples generated.

Image translation technologies map source domain image $X$ to target domain image $Y$ via the generator mapping: $g: X\to Y$. The generator operates by downsampling the $X$ via convolution and pooling operations into latent vector $Z$, then upsampling this latent distribution through upsampling and deconvolution operations. CycleGAN utilizes six loss functions as part of its modeling objective. First, images from domain $X$ is converted into $Y$ via the generator $G$. Here, $L_{GAN}$ is applied to match the distribution of $\hat{Y}=G\left( X \right)$ to $Y$, and utilizes a discriminator for the target domain images, $D_{Y}$:

$$L_{\mathrm{GAN}}\left( X,Y,G,D_{Y} \right)=E_{y\sim p_{r}\left( y \right)}\left[ log\left( D_{Y}\left( y \right) \right) \right]+E_{y\sim p_{g}\left( y \right)}\left[ log\left( 1-D_{Y}\left( G\left( x \right) \right) \right) \right]$$

To maintain structural consistency after translation, the generated image $\hat{Y}$ is backtranslated into the source domain via generator, $F: Y\to X$, and compared to the original source image X via the reconstruction loss function, which is cycle-consistent ($X\to Y \to X$):

$${L_{recon,x}=E}_{x}\left[ \left| \left| F\left( G\left( x \right) \right)-x \right| \right|_{1} \right]$$

Similar operations are applied to the target domain, where $Y$ is mapped to $X$, and a discriminator $D_{X}$ compares $\hat{X}$ to $X via L_{GAN}\left( F,D_{X} \right)$, and back translation to Y for structural consistency and that the recapitulated data resembles its original source:

$${L_{recon,y}=E}_{y}\left[ \left| \left| G\left( F\left( y \right) \right)-y \right| \right|_{1} \right]$$

The cycle-consistent loss is given by:

$$L_{cyc}\left( G,F \right)=\lambda_{a}L_{recon,x}+\lambda_{b}L_{recon,y}$$

Where $\lambda_{a}$ and $\lambda_{b}$ are hyperparameters that place more emphasis on the source versus target domain. Finally, identity losses ensure that if $Y$ were accidentally fed into the generator $G$, it would yield itself via the identity mapping, $Y=G(Y),$ and similarly for $X$, $X=F(X).$ These losses are applied to further regularize the neural network:

$$L_{identity}=\left[ \left| \left| F\left( x \right)-x \right| \right|_{1} \right]{+E}_{y}\left[ \left| \left| G\left( y \right)-y \right| \right|_{1} \right]$$

Together, the final loss for CycleGAN is:

$$L_{CycleGAN}=L_{GAN}\left( G,D_{Y} \right)+L_{GAN}\left( F,D_{X} \right)+L_{cyc}\left( G,F \right)+\lambda_{\mathrm{identity}}L_{identity}\left( G,F \right)$$

Similar *minimax* optimization of the objective with respect to the parameters of $G,F,D_{X},D_{Y}$ yields the ideal generator $G$ for stain conversion that generates realistic samples via the GAN losses and structural consistency via the cycle-consistent reconstruction losses.

**Software Implementation**

Some example scripts for evaluating the CycleGAN model on custom NPY files and automatic conversion and deployment to openseadragon may be found in the following GitHub repository: <https://github.com/jlevy44/HE2Tri> (adapted from <https://github.com/junyanz/pytorch-CycleGAN-and-pix2pix>).

**Supplementary Figure 9:** Screenshots from interactive openseadragon web application to test intermediate iterations of GAN model fitting for model selection; H&E stain is presented side-by-side with converted trichrome stain using the: a) sync view option; b) curtain view option; web application will be further refined for optimal viewing options during prospective tests of clinical deployment
